# Cytosolic localization and *in vitro* assembly of human *de novo* thymidylate synthesis complex

**DOI:** 10.1101/2020.12.23.423904

**Authors:** Sharon Spizzichino, Dalila Boi, Giovanna Boumis, Roberta Lucchi, Francesca Romana Liberati, Davide Capelli, Roberta Montanari, Giorgio Pochetti, Alessio Paone, Serena Rinaldo, Roberto Contestabile, Alessandro Paiardini, Angela Tramonti, Giorgio Giardina, Francesca Cutruzzolà

## Abstract

*De novo* thymidylate synthesis is a crucial pathway for normal and cancer cells. Deoxythymidine monophosphate (dTMP) is synthesized by the combined action of three enzymes: serine hydroxymethyltransferase (SHMT), dihydrofolate reductase (DHFR) and thymidylate synthase (TYMS), the latter two targets of widely used chemotherapeutics such as antifolates and 5-fluorouracil. These proteins translocate to the nucleus after SUMOylation and are suggested to assemble in this compartment into the thymidylate synthesis complex (dTMP-SC). We report the intracellular dynamics of the complex in lung cancer cells by *in situ* proximity ligation assay, showing that it is also detected in the cytoplasm. This result strongly indicates that the role of the dTMP-SC assembly may go beyond dTMP synthesis. We have successfully assembled the dTMP synthesis complex *in vitro*, employing tetrameric SHMT1 and a bifunctional chimeric enzyme comprising human TYMS and DHFR. We show that the SHMT1 tetrameric state is required for efficient complex assembly, indicating that this aggregation state is evolutionary selected in eukaryotes to optimize protein-protein interactions. Lastly, our results on the activity of the complete thymidylate cycle *in vitro*, may provide a useful tool to develop drugs targeting the entire complex instead of the individual components.

## INTRODUCTION

Maintenance of the physiological dNTP levels is critical for genome stability and alteration in their levels have complex consequences [1]. Among *de novo* nucleotide synthesis pathways, deoxythymidine monophosphate (dTMP) synthesis is active in many tissues and is critical for the proliferation of different tumours [2]; it also connects dNTP synthesis with folate and one-carbon metabolism, a complex network of reactions controlling synthesis of a number of different precursors and providing methylation and antioxidant power. All enzymes involved are strictly regulated at the transcriptional and translational level [3–5].

dTMP is synthesized starting from deoxyuridine monophosphate (dUMP) by thymidylate synthase (TYMS: EC:2.1.1.45) in synergy with two other folate-dependent enzymes of the folate cycle: serine hydroxymethyltransferase (SHMT1: EC:2.1.2.1) and dihydrofolate reductase (DHFR: EC:1.5.1.3) [5]. SHMT1 produces 5,10-methylenetetrahydrofolate (CH_2_-THF) from tetrahydrofolate (THF) using serine as one-carbon source. The carbon atom is then transferred from CH_2_-THF to dUMP by TYMS to form dTMP and dihydrofolate (DHF); finally, the NADPH-dependent reduction of DHF to THF, catalysed by DHFR, closes the thymidylate synthesis cycle (Fig 1). dTMP synthesis seems compartmentalized to the mitochondria [6] and the nucleus, where it sustains DNA replication during S-phase or after DNA damage. The three enzymes undergo a SUMO-dependent translocation into the nucleus, where they were shown to assemble in the thymidylate synthesis complex (dTMP-SC), anchored to the lamina by SHMT1, but also present at DNA synthesis sites and interacting with the DNA replication machinery [7–10] (Scheme 1). Complex formation in the nucleus is assumed to be responsible for dTMP synthesis and to prevent genome uracil misincorporation [7,11,12].

**Figure1.**
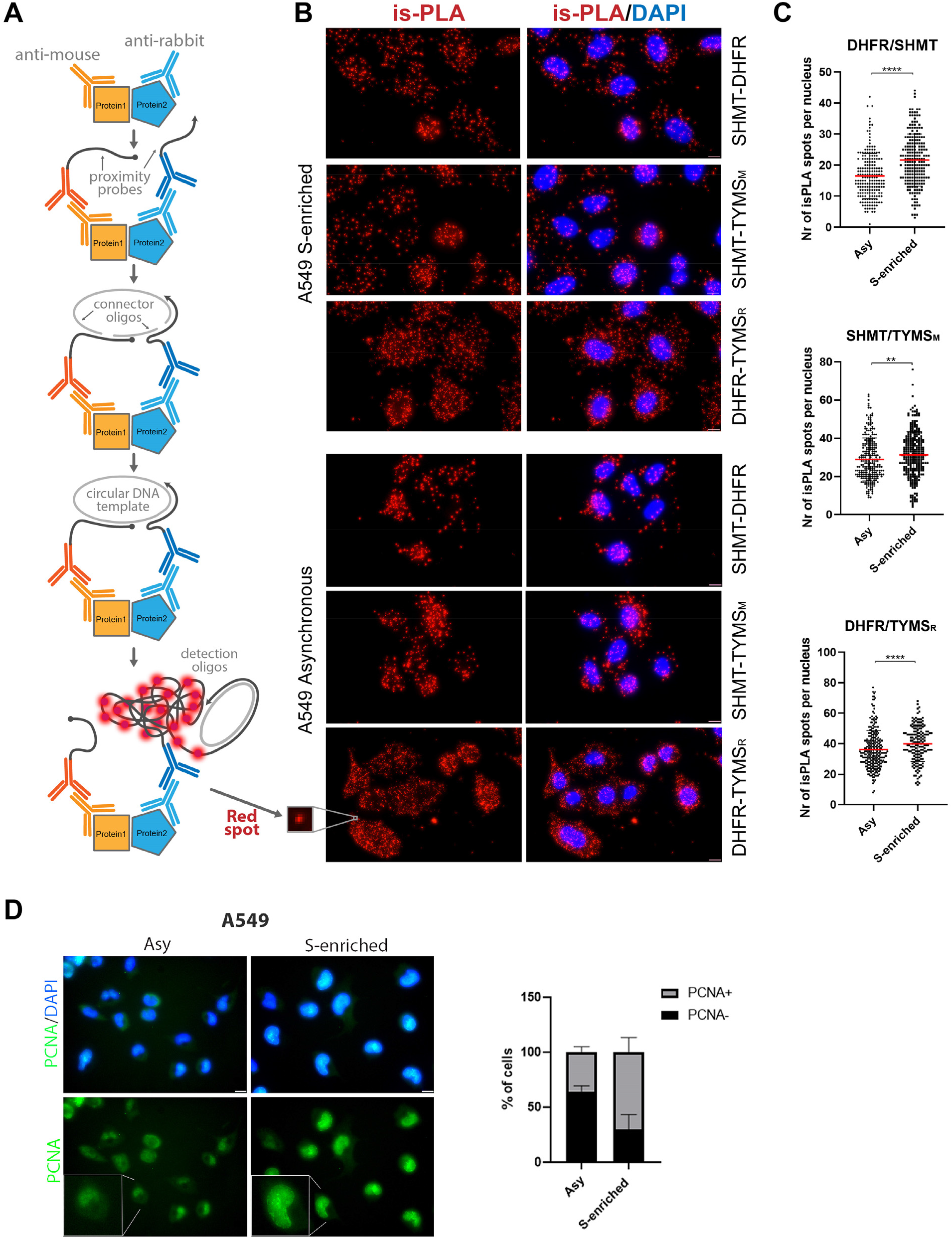
Identification of proteins proximity in cancer cell lines. A) Scheme of the *in situ* Proximity Ligation Assay (is-PLA): 1 - primary antibodies bind specific proteins; 2 - Secondary antibodies conjugated with oligonucleotides (proximity probes) bind to anti-rabbit or mouse primary antibodies; 3 - If the two proteins are interacting (less than 40 nm apart) the proximity probes can hybridize with the two connector oligos; 4 - the ligation step produces a circular DNA template; 5 - Circular DNA is then amplified by DNA polymerase. Detection oligos coupled to fluorochromes hybridize to repeating sequences in the amplicons yielding the is-PLA signal detected by fluorescent microscopy as discrete spots (inset). B) is-PLA signals, corresponding to DHFR/SHMT, SHMT/TYMS and DHFR/TYMS interactions, are shown (red dots). TYMS_M_ and TYMS_R_ both indicate antibodies of TYMS but anti-mouse and anti-rabbit, respectively. The is-PLA spots of interaction of proteins are shown in A549 cells synchronised in the S-phase after a 24 h single thymidine block (upper panel) or in A549 asynchronous cells (bottom panel). Scale bars: 10 μm. The merge with DAPI signal is shown on the right. As a control the assay was repeated using a single antibody directed against SHMT1, DHFR and TYMS (Fig. 3A). C) Nuclear localization of the is-PLA signal (number of spots within the normalized nucleus area) In all the experiments the number of PLA spots increase in the S-enriched cells (T-test: **P < 0.005; ****P < 0.0001). D) On the left: representation of PCNA positive cells, corresponding to S-phase of the cell cycle. Scale bar: 10 μm. On the right: immunofluorescence signal in an asynchronous population or after a single thymidine block of A549. The histogram on the right shows the percentage of PCNA positive or negative cells after and before synchronisation. At least 500 cells were counted per conditions, from three independent experiments, s.d. are shown. After synchronisation, the cells in S-phase are about the 80% of cellular population.

Given their role in cell proliferation, two out of three of the dTMP-SC enzymes - TYMS and DHFR - are targets of widely used chemotherapeutic drugs, such as 5-fluorouracil (5-FU) and antifolates like methotrexate (MTX) or pemetrexed (PTX) [13–15]. In the cells, 5-FU is converted to fluorodeoxyuridine-monophosphate (FdUMP) which covalently binds to TYMS acting as a suicide inhibitor, while the antifolates are cofactor analogues acting as competitive inhibitors [2]. Given the importance of the targeted enzymes in nucleobases metabolism, the side effects of these treatments are severe [16]. Moreover, despite these drugs being in use from the early 60s, the major drawback of this chemotherapy is that cancer cells can rewire their metabolism in response to the lack of THF and dTMP by increasing the expression of both TYMS and DHFR or by upregulating ATP-driven efflux transporters [17–19]. To overcome these issues, a great effort has been made to find new inhibitors targeting other enzymes, such as SHMT [20–25], to date with little results.

An alternative therapeutic strategy would be to target protein-protein interactions (PPI) instead of the single enzymes. Protein complexes and PPI are commonly formed by cells to increase the efficiency, tunability and control over crucial metabolic pathways [26]. Many of these protein assemblies undergo complex dynamics, often controlled by other cellular components such as nucleic acids and lipids and even segregate in membraneless intracellular compartments.

In the present paper, we aimed at understanding how the dTMP-SC complex coherently assembles and functions to ultimately provide the molecular basis to understand how it orchestrates dTMP metabolism. Several features of the complex are still unclear, including how the three enzymes come together, and if the complex is needed to provide dTMP *in situ* during DNA replication/repair or to enhance the catalytic efficiency through a substrate tunnelling mechanism or to generate a binding region to anchor the complex to DNA. Nor it is clear whether the dTMP-SC is formed in the cytosol and needed to control dTTP local pools or may be involved in other functions.

Our results show that the dTMP-SC is abundant in the cytoplasm of both S-phase synchronised and non-synchronised lung cancer cells, suggesting that the interaction between these enzymes may go beyond the nuclear dTMP synthesis and embrace novel regulatory pathways yet to be unveiled. We have successfully assembled the dTMP synthesis complex *in vitro*, employing tetrameric SHMT1 and a bifunctional chimeric enzyme comprising human TYMS and DHFR, in which the two enzymes were still active. To the best of our knowledge, the dTMP-SC has never been isolated before. Only one work reports the interaction of human TYMS and DHFR, which were shown to form a very faint complex *in vitro* [27]. Lastly, we show here that the SHMT1 tetramer is required for optimal complex assembly, suggesting that this aggregation state is evolutionary selected in eukaryotes to optimize PPI, which may indeed represent a promising drug target.

## RESULTS

### *In situ* proximity ligation assay (is-PLA)

To monitor the formation of the dTMP-SC and to obtain novel information on its localization in space and time, we performed an *in-situ* proximity ligation assay (is-PLA). A major benefit of PLA is that it does not require modification or tagging of the proteins of interest, allowing endogenous interactions to be detected with greater sensitivity, with respect to co-Immunopurification methods previously employed for the dTMP-SC [7]. We used both non-synchronised and S-phase synchronised lung cancer cells (A549) to determine the interactions between all the possible couples in the ternary complex: SHMT1-TYMS-DHFR. Is-PLA uses ordinary primary antibodies to detect the proteins of interest, which are then revealed by using secondary antibodies conjugated to DNA oligonucleotides which produce a fluorescent signal in the form of a spot, given by the ligation and amplification of the oligonucleotides that occurs only when the two proteins are interacting or closer than 40 nm (see scheme in Fig. 1A) [28]. As shown in Figure 1B we could detect the positive PLA signals for all the combinations of the three proteins in both synchronous and asynchronous A549 cells. Interestingly, the complex was more abundant in the cytoplasm than in the nucleus, and nuclear localization appears to be enriched in synchronised S-phase cells only to some extent (Fig. 1C).

To confirm the latter observation, we performed a co-localization analysis (Fig 2A) and independently detected the presence of the single proteins (SHMT1, TYMS and DHFR) in the cytoplasm and in the nucleus of both synchronous and asynchronous A549 cells by immunofluorescence (Fig. 2B to D) and by western blot after cell fractioning (Fig. 2F). The three proteins were always present in the cytoplasm, whereas nuclear localization was more abundant in S-enriched cells only for TYMS and SHMT1 (Fig. 2E). This was particularly evident for the latter, suggesting that nuclear translocation of SHMT1 may drive the increase of dTMP-SC formation in the nucleus during the S-phase. Overall, the PLA results suggest that, under these conditions, the dTMP-SC is located in the cytoplasm as well as in the nucleus. This distribution was further confirmed in asynchronous HeLa cells (Fig. 3B).

**Figure 2:**
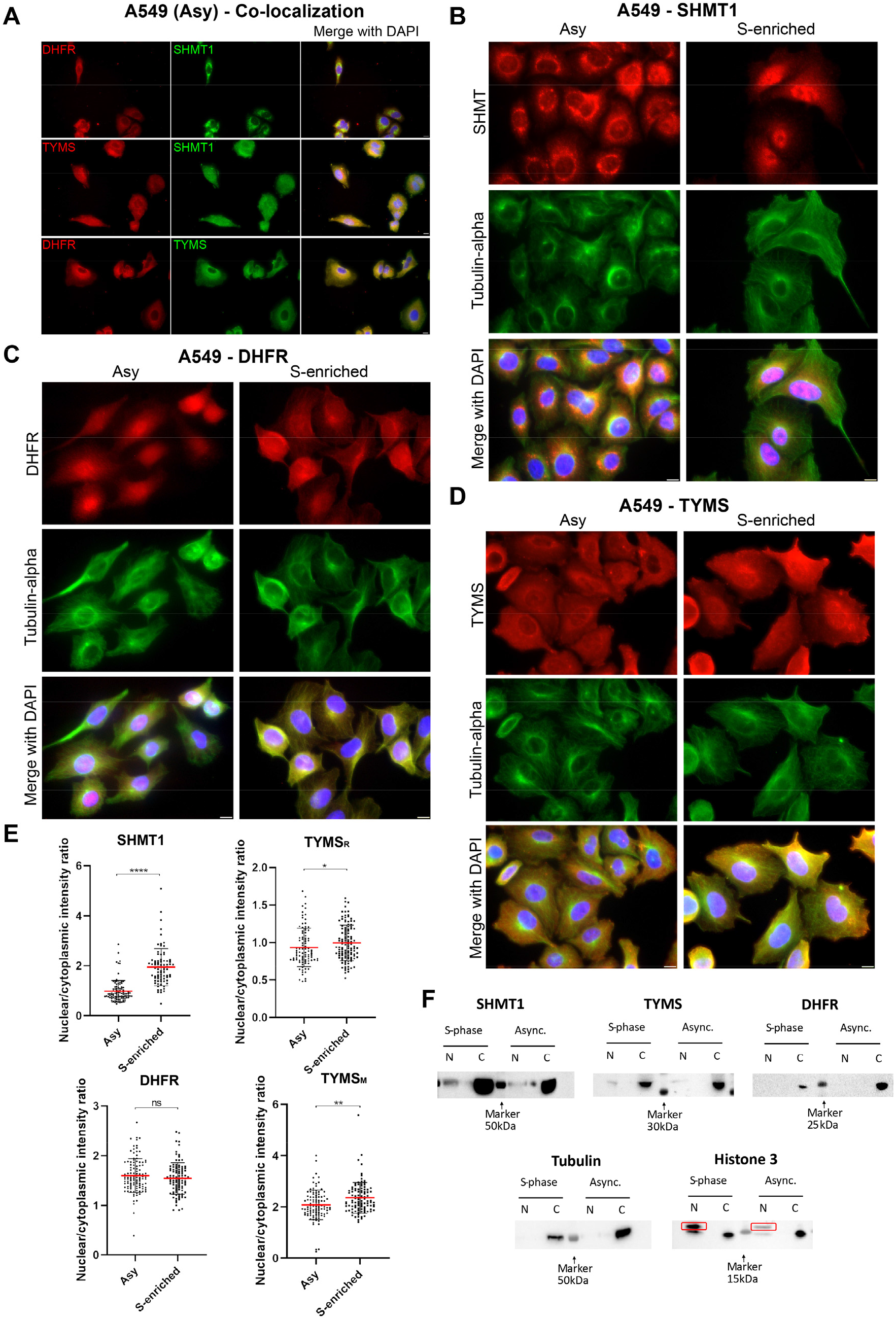
Immunofluorescence analysis. A) Co-localization of DHFR/SHMT, TYMS/SHMT and TYMS/DHFR in asynchronous A549 cells. Co-localization is evident both in the cytosol and in the nucleus for all the three couples. Scale bar: 10 μm. A-B-C) Compartmentalization of the three proteins with respect to the cellular phase by Immunofluorescence analysis. SHMT1 (B), DHFR (C) and TYMS (D) show a cytoplasmatic localization both in the asynchronous and S-phase synchronised cells. Scale bar: 10 μm. E) Nuclear localization (as deduced by the normalized intensity of the fluorescence signal) is clearly more abundant in the S-enriched cells for SHMT, a slight increase is observed for TYMS (with both the antibodies used in the PLA experiments; rabbit - TYMS_R_ and mouse – TYMS_M_. See experimental section). No change was observed for DHFR. (T-test: *P < 0.05; **P < 0.005; ****P < 0.0001). F) Western blot of subcellular fractionation of both asynchronous and S-phase enriched A549 cell lines. WB analysis was performed using a 1:1000 dilution of the primary antibodies. For Histone H3 the correct band is boxed in red. The other bands detected in the cytosol, have a molecular weight lower than 15kDa and are also present in the nuclear fractions. Giving that the Mw of Histone H3 is 17kDa, and the bands recur in all the four lanes, it is plausible that they are detected because of non-specific interactions of the primary or secondary antibody. In this experiment the quantitative analysis was not performed, but it is still possible to detect the bands of SHMT1 and TYMS only in the nucleus of the S-phase enriched cells. As detected with the IF experiments, it was not possible to perceive an increase of the presence of DHFR in the nucleus of the synchronised cells.

**Figure 3:**
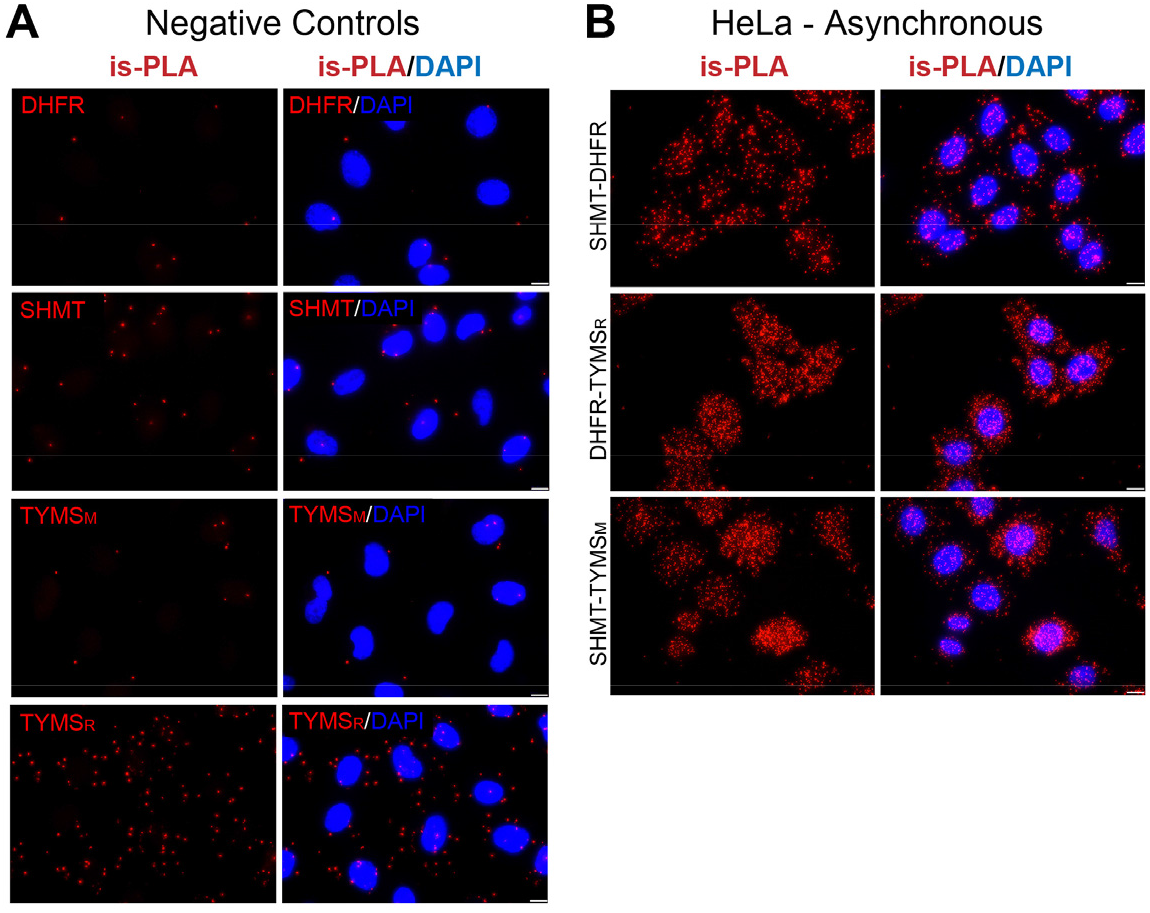
is-PLA negative controls and on HeLa cell. A) Negative controls of is-PLA signals. PLA experiment was performed without one of the primary antibodies. In these conditions there is no or little PLA signal respect to that observed with both primary antibodies. TYMS_M_ and TYMS_R_ both indicate antibodies of TYMS but antimouse and anti-rabbit, respectively. B) is-PLA signal of interaction is shown in HeLa non-synchronized cells. In A and B panels scale bar: 10 μm.

These results indicate that the three enzymes are able to assemble the dTMP-SC complex even in the absence of DNA or lamina proteins as binding partners [7], suggesting that it is possible to assemble the dTMP-SC *in vitro* starting from the purified proteins. Our strategy to achieve this goal was to obtain a binary interaction, by assembling SHMT1 with a TYMS-DHFR fusion protein.

### Rational design of the chimeric bi-functional protein

To understand how DHFR and TYMS interact with each other and to design a chimeric fusion protein (hereinafter called Chimera), bifunctional enzymes endogenously expressed in several Protozoa have been taken as an example. In particular, two classes of DHFR-TYMS bifunctional enzymes can be distinguished, which differ mainly in the length of the linker between the two domains and consequently in the orientation with which the DHFR domain contacts the TYMS domain [29]. The crystallographic structure of DHFR-TYMS from *Trypanosoma cruzi* (PDB ID: 2H2Q) [30] was chosen as a representative for the short linker class, while the structure from *Babesia bovis* (PDB ID: 3I3R)[31] was selected among several DHFR-TYMS structures with long linkers from different organisms, due to the higher identity with the human DHFR sequence (33% and 55% similarity, respectively). The crystallographic structures of human DHFR and TYMS were aligned to the two representatives of the bifunctional enzymes to identify the two possible interaction interfaces. An analysis of the evolutionarily conserved residues at the interfaces showed that in both cases none of the residues involved in the stabilisation of the DHFR-TYMS complex was conserved. Following this evidence, the surface potential at the interface was evaluated and a perfect complementarity of the surface charge of human proteins was revealed at the level of the interaction interfaces originating after a structural alignment with *B. bovis* DHFR-TYMS (Fig. 4A-B). This suggests that the contacts between DHFR and TYMS are nonspecific and that the interaction interface on TYMS provides only an attractive force for DHFR and not an orienting one, as previously suggested [29].

**Figure 4:**
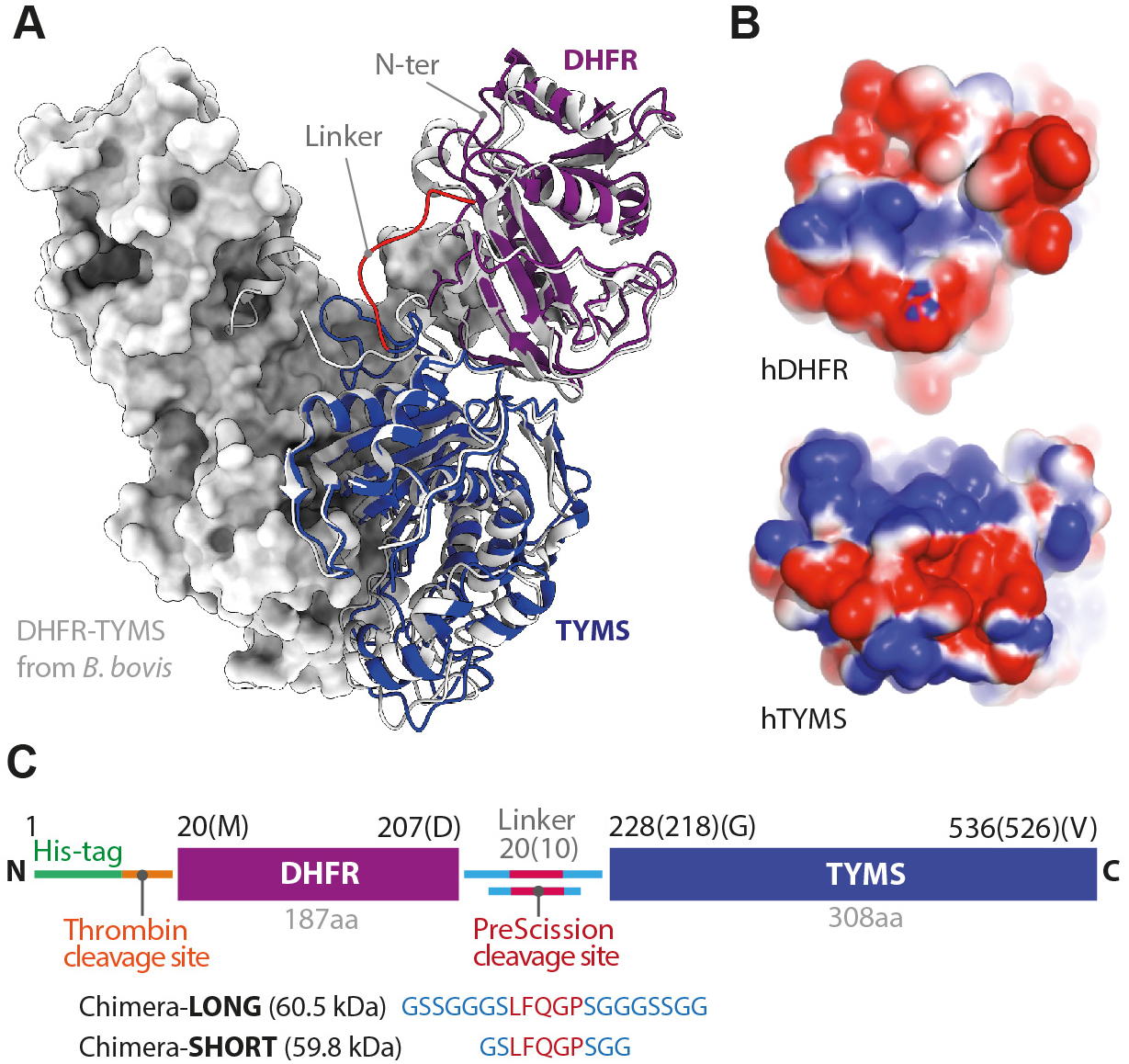
Designing human DHFR-TYMS Chimera. A) Cartoon representation of the chimeric model of human DHFR-TYMS (in *purple* and *blue* respectively; the linker is shown in *red*; the C-terminal position is also indicated) superposed with the structure of the bifunctional enzyme from *B. bovis* (PDB ID: 3I3R [31]) in *light grey*. The enzyme is dimeric, the partner subunit is shown as surface representation. B) Electrostatic surface potentials - calculated using APBS (Adaptive Poisson-Boltzmann Solver) [57] at the modelled interface between hDHFR and hTYMS, showing a perfect complementarity. Partially positive or negative regions are indicated in blue and red, respectively. C) Scheme of the final constructs differing only for the linker length and consequently named *Chimera-Long* and *Chimera-Short*.

Once the interaction modality was chosen, the following step was to design the linker. Even though in the selected interaction mode the C-terminus of DHFR and the N-terminus of TYMS are spatially near to each other, a knot would be created by joining them directly, which could prevent the Chimera fusion protein from folding correctly. For this reason, two different linkers were designed, a short 10 aa linker and a long 20 aa one. The linkers were designed to be flexible but with a higher content of Ser residues compared to the canonical (GGGGS)n flexible module [32], because bioinformatic analysis showed that the linker would be completely exposed to the solvent. Moreover, a sequence sensitive to digestion by PreScission protease was introduced to allow the separation of the two enzymes *in vitro* when needed. The two final linkers were GSSGGGSLFQGPSGGGSSGG for Chimera-Long and GSLFQGPSGG for Chimera-Short (Fig. 4C). The 3D structure of the designed bifunctional enzymes was then modelled based on homology using as templates the structures of hDHFR and hTYMS separated but aligned with the bifunctional enzyme from *B. bovis*. Finally, we used the crystallographic structure of TYMS from *Mus musculus*, to model the N-terminal residues of hTYMS, which is flexible and does not appear in any of the crystallographic structures from *Homo sapiens* available in the PDB.

### Expression, purification, biochemical and biophysical characterization of the chimeric constructs

Both constructs were expressed with an N-terminal histidine-tag and were purified by immobilized metal affinity chromatography (IMAC), eluting between 150 mM and 200 mM imidazole. The constructs showed to undergo proteolysis in the linker region, as highlighted by the low molecular weight species detected by SDS PAGE and western blot (WB) of the purified fractions (Fig. 5A). This phenomenon was more evident at pH higher than 7.5 and was more pronounced for Chimera-Short. To remove the proteolyzed protein, the fractions containing the two constructs were concentrated and further purified by Size Exclusion Chromatography (SEC). Both Chimera-Long and Short eluted as dimers (Fig. 5B), suggesting that at least TYMS was correctly folded in the Chimera, as human TYMS is dimeric. After purification, no further proteolysis occurred, and the proteins were stable for at least one week at 4 °C (Fig. 5C). The UV spectra of Chimera showed a shoulder around 325 nm, likely due to NADPH bound to DHFR (Fig. 5D). This result suggested that also DHFR was correctly folded in the Chimera. Analysis by CD spectroscopy confirmed that both Chimera-Short and Long are folded and stable up to 45° C, with an apparent melting Temperature (T_m_) of 53.5° C (Fig. 5E-F)

**Figure 5:**
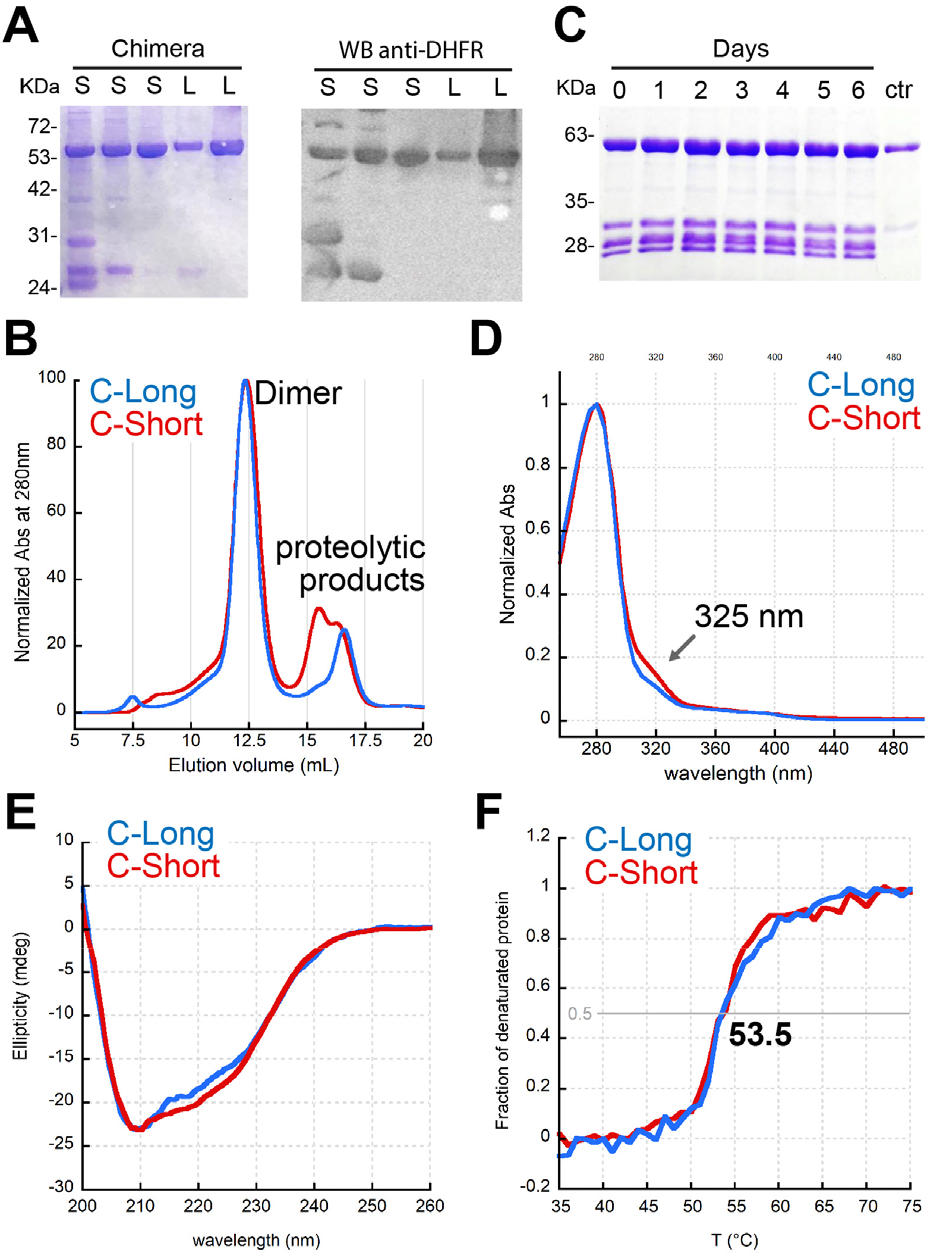
Purification and spectroscopic characterization of Chimera. A) SDS-PAGE and WB of IMAC elution peaks (150-200mM imidazole) of different preparations of Chimera Short (S) and Long (L). B) Typical SEC chromatogram of IMAC fractions of Chimera Constructs. The proteolyzed proteins are well separated from the main peak, containing the full-length protein eluting as a dimer. The signal is normalized for a better comparison (column: superdex 200 10/300). C) Auto proteolysis assay: after IMAC, a small amount of Chimera Short was kept at 4°C for 6d, to test whether the purified protein undergoes proteolysis in the purification buffer. Ctr = protein after SEC. No further proteolysis was observed. Typical final yields were between 6 and 4 mg/L of culture for Chimera-Long and Short, respectively. D) Normalised UV spectra of Chimera-constructs in 100 mM potassium phosphate pH 7.4. The shoulder at 325 nm is likely due to bound NADPH. E) Dichroic spectra of 10 μM Chimera constructs at 20 °C in a 1 mm quartz cuvette. F) Normalized thermal denaturation profiles of 10 μM Chimera constructs.

### Activity of Chimera and of the full thymidylate cycle

As a final quality control, the complete thymidylate cycle was analysed to assess the full functionality of the fused enzymes, as described in Figure 6. In a first assay, the reductive methylation of dUMP to dTMP catalysed by TYMS was assessed by following the increasing absorbance at 340 nm due to the formation of DHF from CH_2_-THF (reaction 1 - Fig. 6A scheme). By increasing the enzyme concentration at constant substrates concentrations, a linear increase of TYMS activity is observed for both Chimera-Short and Long (Fig. 6B). However, since DHFR is fused to the N-terminal of TYMS, and mutation in this region are known to affect catalysis [33,34], as a control, we also performed a complete characterization of TYMS activity that yielded kinetic parameters (K_cat_ = 0.5 s^−1^; K_m_ for CH_2_-THF = 2.9 ± 0.5 μM; K_m_ for dUMP = 7.1 ± 1.0 μM) very similar to those reported for wild type TYMS [35] (Fig 6C), indicating that TYMS is correctly folded and fully functional in the designed constructs.

**Figure 6:**
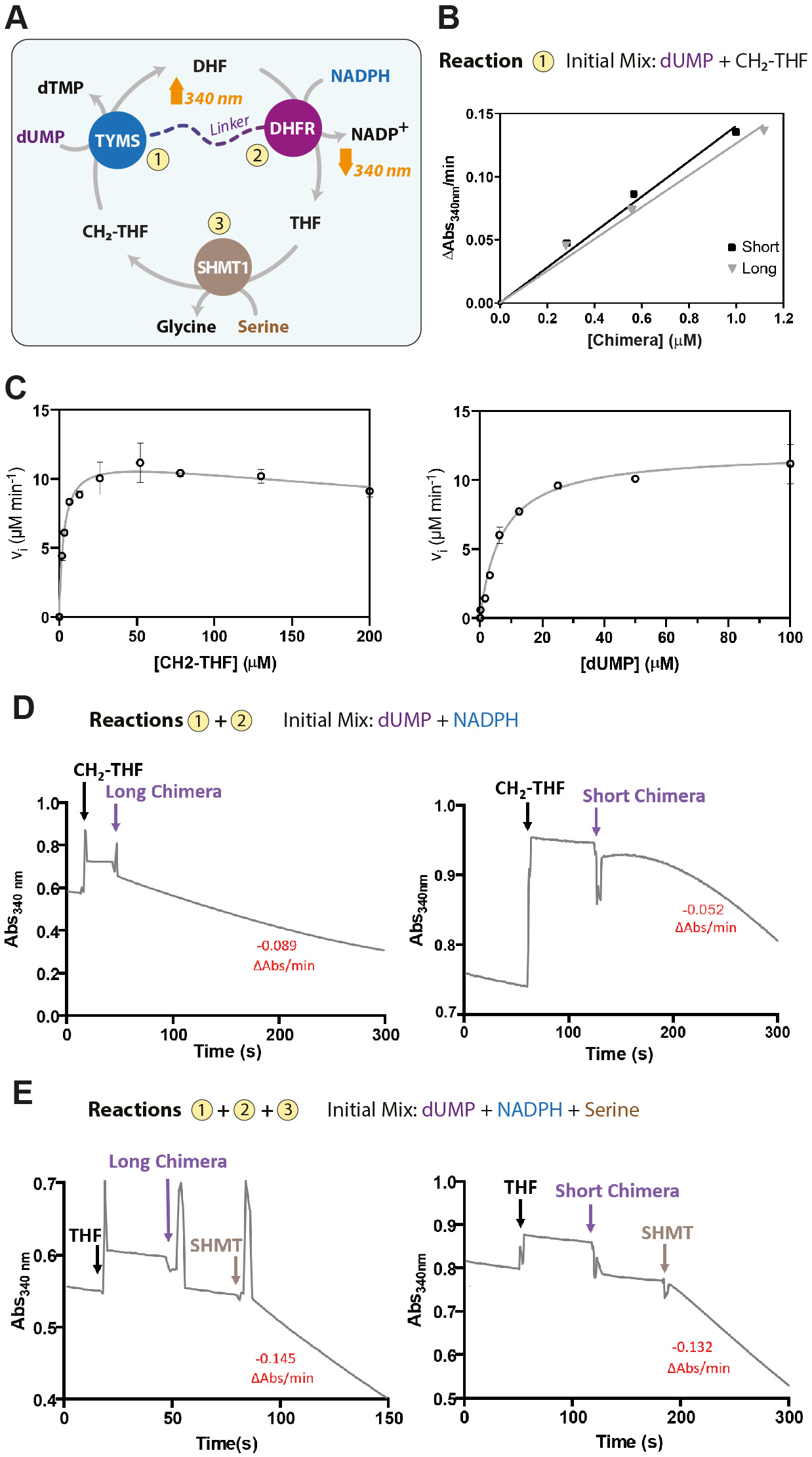
Activity assays: restoring the thymidylate cycle *in vitro*. A) Scheme of the reactions assayed to test the catalytic activity of Chimera. All the reactions were performed at 20°C in 20 mM K-Phosphate pH 7.2, 75 mM β-Mercaptoethanol. B) Reaction 1; plot of the initial rate of dTMP formation as a function of Chimera concentration at constant substrate (0.1 mM dUMP, 0.2 mM CH_2_-THF); Chimera-Short (*black squares*), Chimera-Long (*gray triangles*). C) Plot of initial rates of reaction 1 as a function of CH_2_-THF concentration (left panel; 0.1 mM dUMP fixed concentration) and of dUMP concentration (right panel; 0.05 mM CH_2_-THF fixed concentration). D) Reactions 1 + 2; time course of the coupled reactions of TMYS and DHRF. The observed rates (ΔAbs@340 nm/min) are also reported in red. To 0.1 mM dUMP and 0.1 mM NADPH, was sequentially added 0.1 mM CH_2_-THF and 0.5 μM Chimera-Long (*left*) or Chimera-Short (*right*). E) Reactions 1 + 2 + 3; to the reaction mixture containing 0.1 mM dUMP, 10 mM serine and 0.1 mM NADPH, were added 16 μM THF, 0.5 μM Chimera-Long (*left*) or Chimera-Short (*right*) and finally 0.5 μM SHMT1. The thymidylate cycle can only start after the addition of SHMT1 that is needed to convert THF to CH_2_-THF.

Then, the DHFR activity of Chimera was investigated in a coupled assay, in which the DHF produced by TYMS activity is then reduced to THF by DHFR, with NADPH oxidation (reactions 1 and 2 – Fig. 6A scheme). The reaction is observed by following the decrease of absorbance at 340 nm due to NADPH oxidation. The time course of the reaction steps is shown in Fig. 6D. With this experimental setup both constructs were able to catalyse the coupled reactions at similar rates, indicating that also DHFR is correctly folded and functional. Finally, the complete thymidylate cycle was then tested. The initial assay mix contained dUMP, serine and NADPH; then THF, Chimera and finally SHMT1 were added. In this experiment, shown in figure 6D, the Chimera activity can start only when the substrate CH_2_-THF is produced by SHMT1, the reaction proceeds until dUMP and NADPH are consumed, while the folate species cycles, thus reconstituting the functionality of the thymidylate cycle *in vitro*.

### *In vitro* analysis of the dTMP synthesis complex

The formation of dTMP-SC between Chimera and SHMT1 was initially investigated by far-western blotting (FWB) and immunopurification (IP). The dissociation constant was estimated by surface plasmon resonance (SPR) and the features of the complex were further explored by analytical gel filtration. Finally, the effect of substrates on complex formation was examined by differential scanning fluorimetry (DSF).

#### Far Western Blotting (FWB) and Immunopurification assay (IP)

Formation of the complex between SHMT1 and the Chimera construct was initially evaluated by FWB assay. In this experiment, the purified Chimera constructs and SHMT1 were resolved by SDS-PAGE and electro-blotted on a PVDF membrane. The proteins were then renatured directly on the membrane and incubated with the bait protein (either SHMT1 or Chimera). In this way, Chimera and SHMT1 can form a complex in native conditions on the membrane that can be detected by specific antibodies (scheme in Fig. 7A). As shown in figure 7A, the antibodies directed against DHFR and SHMT1 detected the presence of Chimera-Long in the lane of SHMT1 and *vice versa*. These results indicate that Chimera-Long and SHMT1 interact with each other, but not with the control protein. The same results were not observed when using the Chimera-Short construct (Fig. 7B). This result suggests that the shorter linker might prevent the optimal orientation of TYMS and DHFR with respect to SHMT1. To further investigate the stoichiometry of the dTMP-SC complex, the FWB was repeated using a Chimera-Long and a dimeric variant of SHMT1 (H135N-R137A) [36]. The inability of dimeric SHMT1 to form the complex under the same experimental conditions (Fig 7C), indicates that the tetrameric state of SHMT1 is crucial for complex formation.

**Figure 7:**
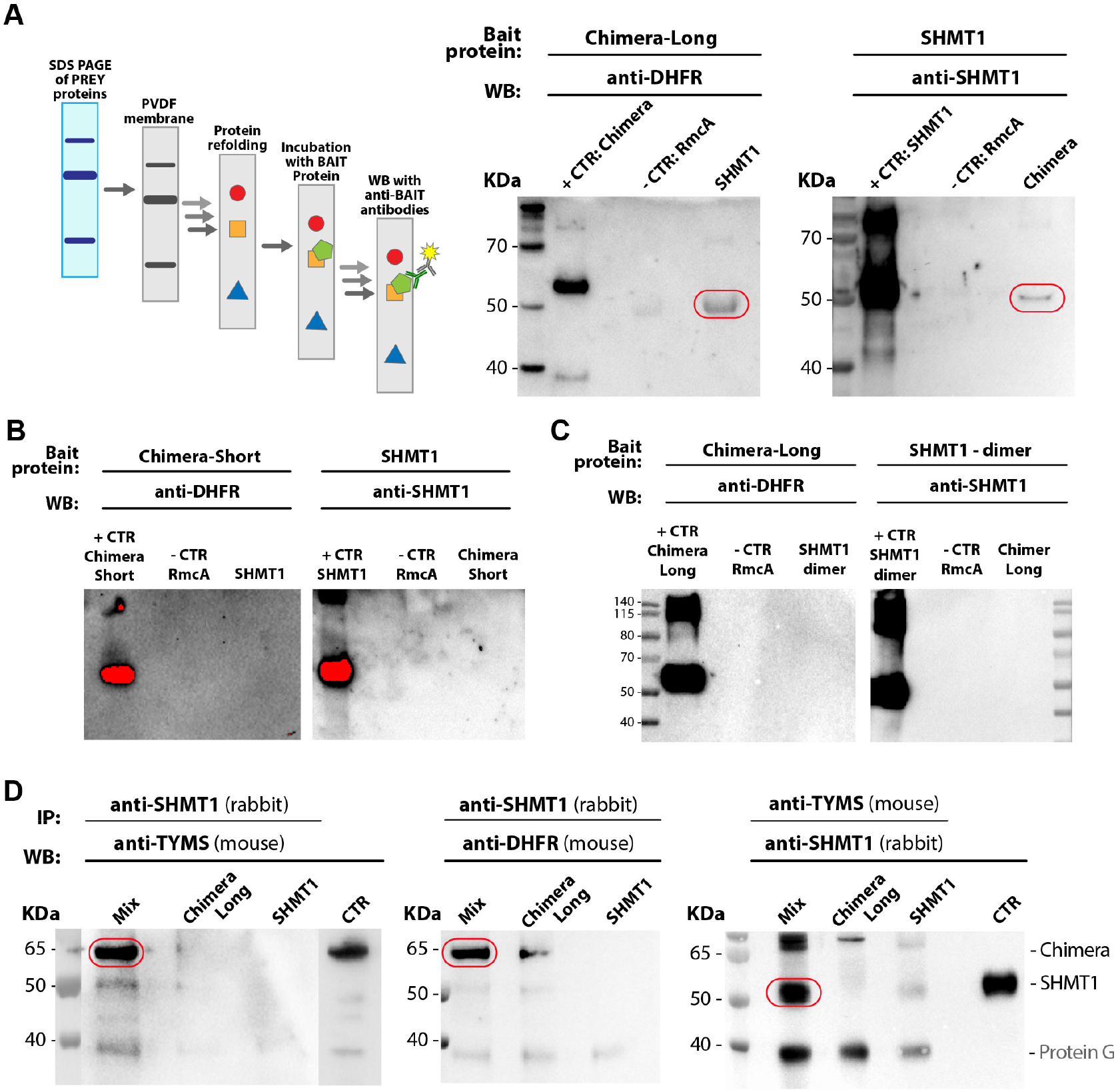
Immunopurification assay (IP) and Far Western Blotting (FWB). A) FWB: The prey proteins (SHMT1 or Chimera-Long) were resolved alone by SDS-PAGE, electro-blotted on a PVDF membrane, refolded and then incubated with 50 ng/ml of the bait protein at 4°C o.n. In the left image the band detected at SHMT1 lane and height represents Chimera, and the band detected at Chimera lane and height, on the right image represents SHM1. RmcA (a bi-domain bacterial protein construct from *P. aeruginosa* whose two domains have a molecular weight comparable to DHFR and TYMS) was used as a control [59]. The formation of the complex between SHMT1 and Chimera-Short (panel C) or SHMT1 dimeric mutant and Chimera-Long (panel D) was tested by following the experimental set-up previously described. Nevertheless, and despite over exposition of the membranes, in both cases formation of the complex was not observed. D) IP experiment: SHMT1 and Chimera-Long were mixed in a 1:2 ratio at a final concentration of 18 μM for SHMT1 and 36 μM for Chimera-Long. The proteins were attached to the Protein G-agarose beads by using specific antibodies (anti-TYMS or anti-SHMT1). The samples were incubated at 4°C o.n. In the first two images, on the left, the detected band represents Chimera-Long whereas in the right image the detected band refers to SHMT1.

The assembly of the dTMP-SC super-complex was confirmed by IP assay, which showed that SHMT1 and Chimera-Long are interacting *in vitro* (Fig. 7D). The proteins were bound to the protein G-agarose beads by using either anti-SHMT1 or anti-TYMS antibodies; Chimera and SHMT1 were only present in the SHMT1 plus Chimera mixture (MIX sample), when detecting respectively with anti-DHFR or anti-TYMS and with anti-SHMT1 antibodies, but not in the individual SHMT1 or Chimera samples (Fig. 7D).

Due to these results and to the higher degree of purity and less tendency to proteolysis of the Chimera-Long construct, we chose to proceed using only this construct for further analysis of complex formation. Here and after, if not otherwise specified, we will refer to the Long construct as Chimera.

#### Surface Plasmon Resonance (SPR)

After evidence of complex formation, we performed SPR experiments to evaluate the binding affinity. We chose to immobilize Chimera and to use tetrameric SHMT1 as analyte. Even though the precise stoichiometry of the putative complex is unknown, we previously showed that the dimeric SHMT1 variant is unable to form the complex. In addition, the higher symmetry of SHMT1 suggests that two Chimera dimers (DFHR-TYMS:TYMS-DHFR) may bind to one SHMT1 tetramer. In this case, the tetrameric form of SHMT1 would have two identical binding sites. This would allow the analyte to bind first with one site to the ligand Chimera and then to bind to another Chimera molecule that is in close contact to the second ligand site. The second binding will give a stabilization of the ligand-analyte complex without extra response but shifts the equilibrium constant to a more stable interaction. This effect, which is often referred to as *avidity*, was observed in the case of SHMT1 binding to Chimera, suggesting that indeed SHMT1 may bind more than one Chimera dimer. A bivalent analyte gives rise to two sets of rate constants, one for each binding step, but the meaning of the two sets of rate constants and particularly the second set is very difficult to interpret. However, the avidity effects can be reduced by using very low ligand levels and high analyte concentrations. Low ligand levels give sparsely distributed ligand on the chip, with less chance of two ligand molecules being within reach of a single analyte. On the other hand, high analyte concentration competes out the second binding site, favouring the formation of 1:1 complex. For this reason, we used a very low immobilization density of the ligand (175 RU on Ch 3) and high analyte concentrations (ranging from 85 to 2.7 μM). In this way, the 1:1 bimolecular interaction model gave a reasonable fit of the experimental curve (that is clearly biphasic due to the residual avidity effect) with an estimated K_d_ of 2.5 and 2.9 μM on Channel 1 and 3 of the chip, respectively. Moreover, to exclude mass-transfer we replicated the same experiment at two different flow rates (75 μl/min and 150 μl/min) obtaining substantially the same value of K_d_ (in the presence of mass transfer limitation we should have obtained different K_d_’s). This data show that the interaction between Chimera and SHMT1 is specific and that the dissociation constant is in the low μM range (Fig. 8).

**Figure 8:**
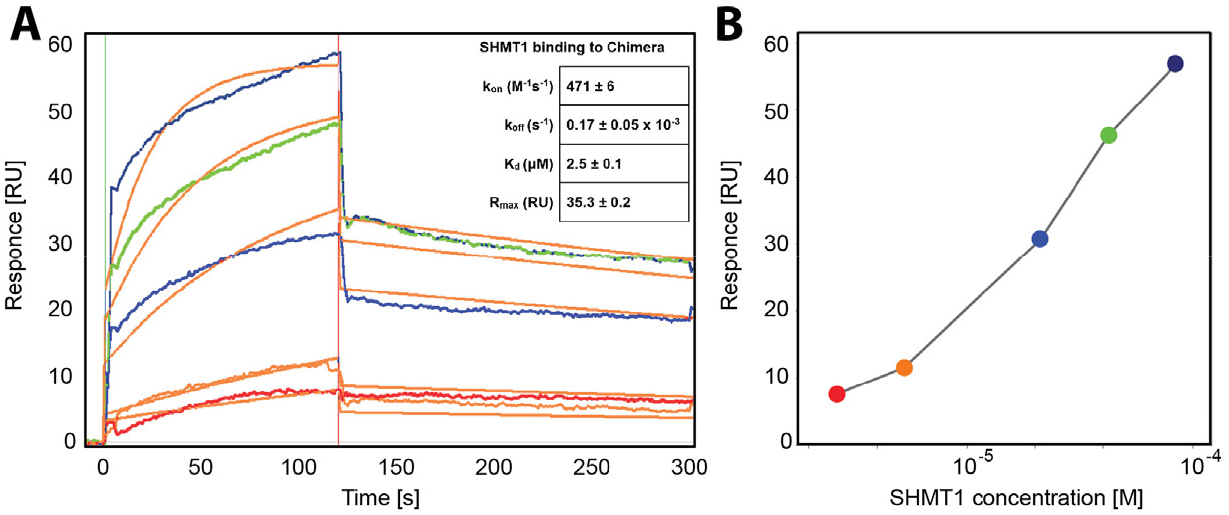
SPR experiment. A) Full kinetic analysis of SHMT1 (85, 42.5, 21.2, 5.3 and 2.66 μM) binding to Chimera (channel 3). Orange lines represent the global fits of the data to a 1:1 bimolecular interaction model. The calculated kinetic parameters - affinity (K_d_) and rate constants (k_on_, k_off_) - for the SHMT1-Chimera interaction are also displayed. B) Dose response plot of the interaction between SHMT1 and Chimera at different concentrations of SHMT1 (85.0, 42.5, 21.2, 5.3 and 2.7 μM) showing a linear dependence.

*Analytical gel filtration* experiments were then carried out to isolate the complex after mixing SHMT1 and Chimera in three different molar ratios, keeping protein concentration above 30 μM, i.e. ten-fold higher than K_d_ obtained by SPR. Unfortunately, although we used an analytical HPLC column, it was not possible to resolve the peaks of the single proteins and of the putative complex. Both Chimera (in the dimeric form) and SHMT1 (tetramer) have similar hydrodynamic radii and eluted between 6.7 and 6.9 ml (Fig. 9A). Chimera also showed a peak at 6.2 ml, likely corresponding to a higher oligomeric state, as TYMS was recently shown to form octamers depending on protein concentration [37]. The signal relative to this last species is expected to cover the one of the complexes. However, with respect to the single proteins, a slight decrease in elution volumes was observed when the samples containing both proteins were injected. The difference can be qualitatively highlighted by subtracting the signal of the individual proteins to these chromatograms. Therefore, we carefully collected the different fractions corresponding to these regions, where SHMT1 should elute only if in complex with Chimera. Western blot analysis showed that SHMT1 was indeed present in the fraction eluting at higher molecular weights, likely corresponding to the complex (Fig. 9). This is particularly evident in the SHMT1:Chimera - 1:2 molar ratio sample (Fig. 9B).

**Figure 9:**
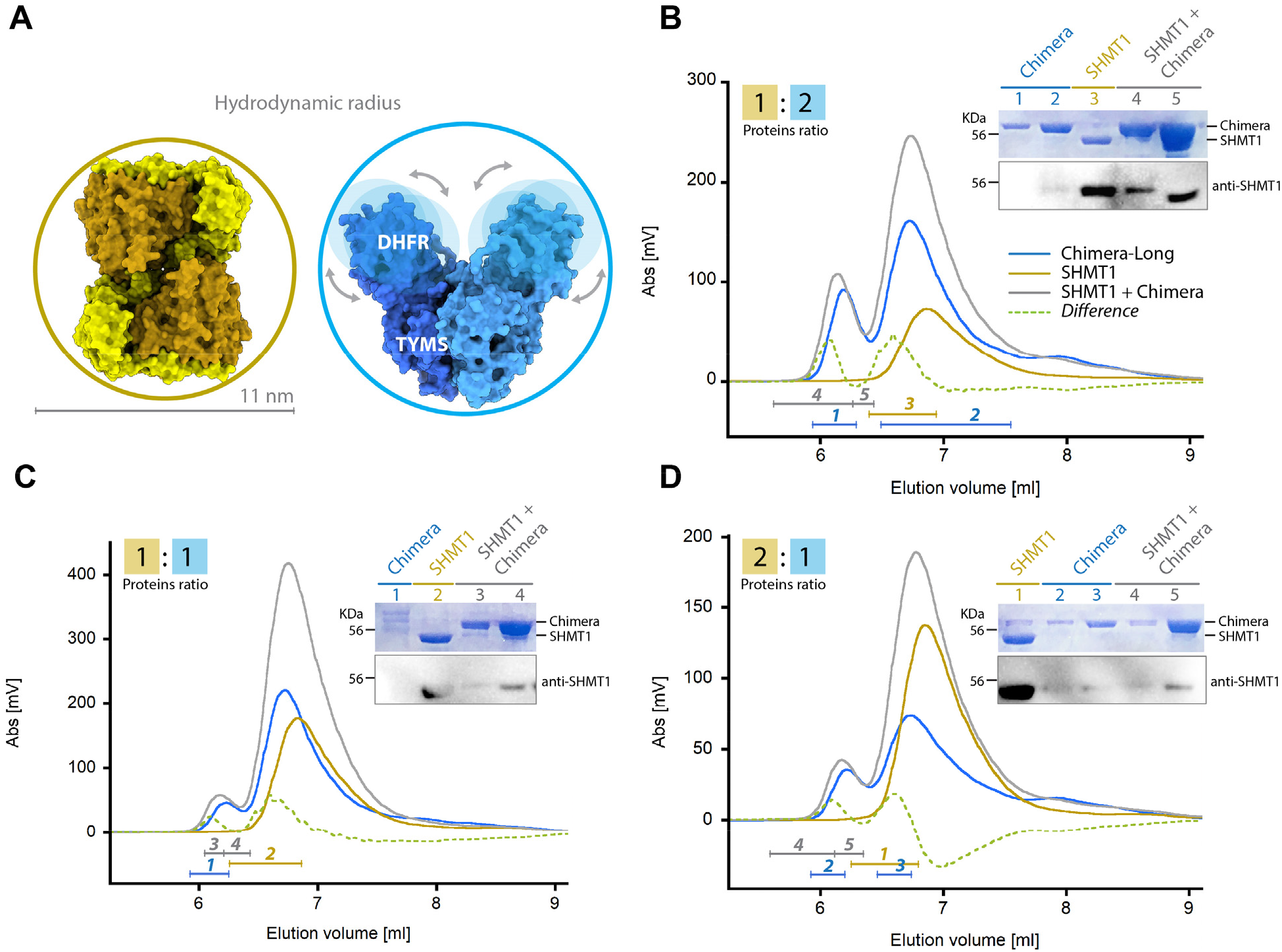
Size exclusion chromatography. SHMT1 and Chimera were mixed at three different molar ratios. 100 μl of protein solutions were injected in a G3000PWXL analytical gel filtration column and detected by following the absorbance signal at 280 nm. A) *On the left*: The calculated hydrodynamic diameter of the SHMT1 tetramer (212 kDa), as extrapolated from the crystal structure (PDB 1BJ4 [64]), is 10.9 nm and SHMT1 elutes at 6.9 ml corresponding to 10.1 nm with respect to the calibration curve of the column. *On the right*: model of dimeric Chimera (this work); Chimera displayed a slightly larger diameter, eluting at 6.7 ml, corresponding to 10.7 nm. This is likely due to the relative mobility of the DHFR domain with respect to the TYMS. Chimera also shows a higher Mw species peaking at 6.2 ml (12.1 nm - 625 kDa). B) Chromatogram of SHMT1+Chimera mixed in a 1:2 ratio (*gray*). Control samples, SHMT1 and Chimera alone, are shown in *brown* and *blue*, respectively. The difference between the chromatogram of the mixed proteins and the controls is shown as a *dashed green* line, the two difference peaks correspond to 12.7 nm - 875 kDa (6.0 ml) and 11 nm - 320 kDa (6.6 ml), respectively. The analysed fractions are indicated below the traces in the same colour of the sample they refer to and numbered according to the loading position in the SDS-PAGE and WB (anti-SHMT1). Panel C) and D) Same as panel B but with different molar ratios (indicated in the top left corner).

#### Effect of substrates and ligands on dTMP-SC formation

The effect of substrate/ligands on *in vitro* complex formation was assessed using DSF. Purified SHMT1 and Chimera were mixed and samples incubated o.n.. SHMT1 alone shows a melting temperature (T_m_) of 57.1±0.1 °C, whereas the T_m_ of Chimera is 48.5±0.5 °C (Table 1). When the two proteins were mixed, the observed T_m_ was 50.2±0.1 °C (Fig. 10A). The change of the denaturation profile is likely due to the interaction taking place between SHMT1 and Chimera. The effect of the presence of substrates, such as 5-formyl-tetrahydrofolate (CHO-THF), dUMP, and NADPH was also tested. While CHO-THF and NADPH had no significant effect, dUMP stabilized Chimera and therefore the Chimera-SHMT1 complex, leaving the T_m_ of SHMT1 unaffected (Fig. 10B and Table 1). Moreover, the slope of the transition increases for the complex indicating that the denaturation process is more cooperative in the presence of dUMP. To confirm that the change in the denaturation profile observed in the presence of dUMP is due to complex formation we performed the same experiment using the Chimera-Short construct that fails to form the complex. In this case the presence of dUMP while stabilizing Chimera-Short (which as expected displayed an overall lower stability), has no significant effect on the stability of the mixed sample (Fig. 10C).

**Figure 10:**
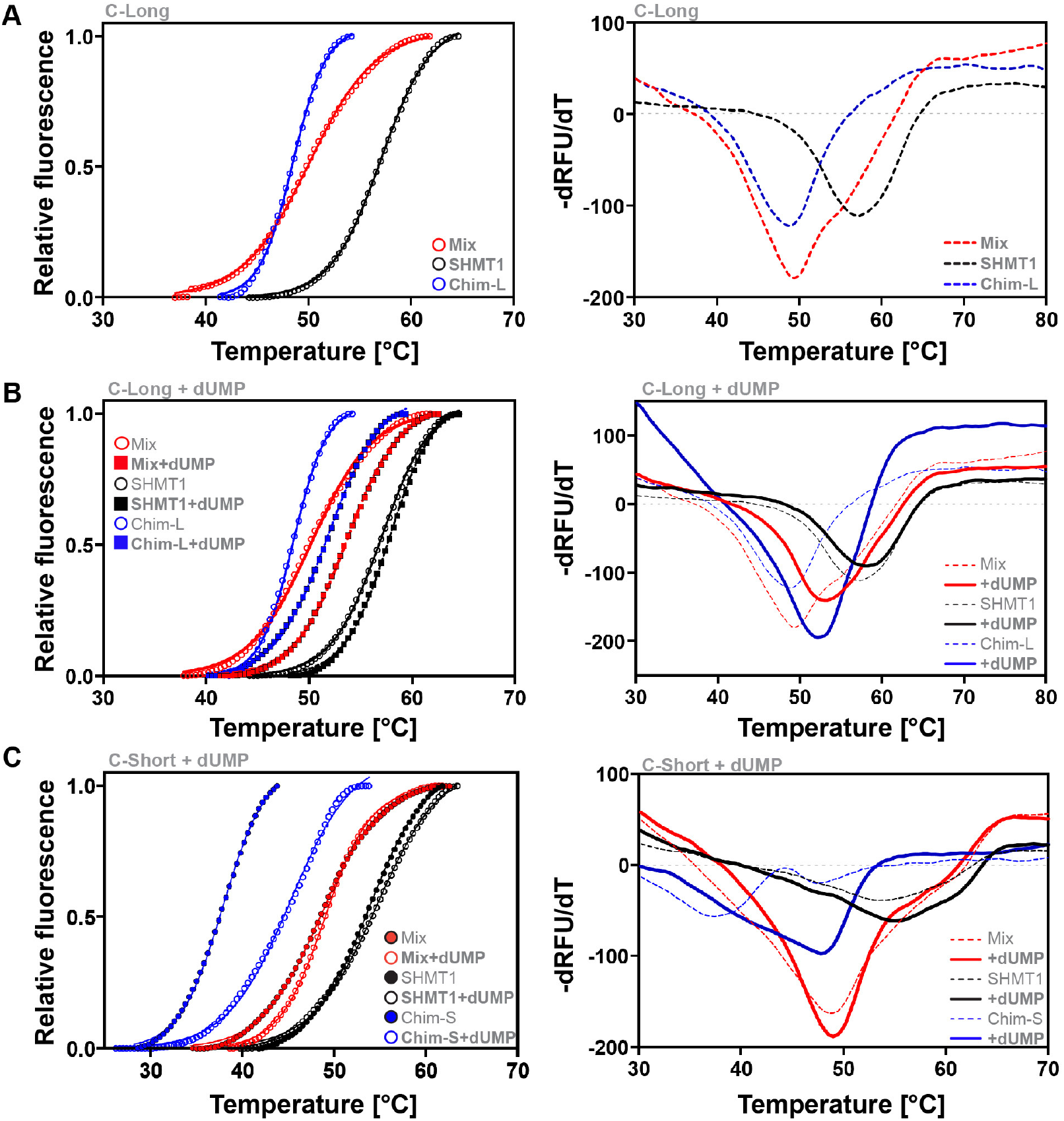
Differential scanning fluorimetry. SHMT1 and Chimera constructs were mixed at a final concentration of 0.5 μM. When present, the concentration of CHO-THF and NADPH were 0.1 mM, dUMP was 1 mM. All samples were incubated o.n. at 4 °C in 20 mM Na Hepes - pH 7.5 and 50 mM NaCl, before starting the analysis. A) denaturation profiles of SHMT1, Chimera-Long and of the mixed proteins in the absence of ligands or substrates. B) Denaturation profiles of SHMT1 and Chimera-Long in the presence of dUMP (square markers and continuous lines) compared to the signals obtained with no ligands/substrates (circle markers and dashed lines). C) Same profiles as in panel B with the Short instead of Long construct. For panels all panels, plots in the left column show the change in fluorescence as a function of temperature, while in the right column the denaturation profiles are plotted as the first derivative of the fluorescence emission as a function of temperature. The calculated T_m_ for all the substrates assayed are reported in Table 1.

**Table 1.** Substrate effect on complex thermal stability: T_m_ (°C)

|  | **SHMT1** | **Chimera-Long** | **Mix** |
| --- | --- | --- | --- |
| **Buffer** | $57.1 \pm 0.1$ | $48.5 \pm 0.5$ | $50.2 \pm 0.1$ |
| **CHO-THF** | $57.7 \pm 0.2$ | $48.2 \pm 0.5$ | $50.5 \pm 0.2$ |
| **NADPH** | $57.3 \pm 0.5$ | $48.3 \pm 0.1$ | $50.7 \pm 0.1$ |
| **dUMP** | $57.8 \pm 0.7$ | $51.8 \pm 0.3$ | $53.6 \pm 0.1$ |
|  | **SHMT1** | **Chimera Short** | **Mix** |
| **Buffer** | $53.8 \pm 0.7$ | $37.9 \pm 0.2$ | $48.5 \pm 0.2$ |
| **CHO-THF** | $54.0 \pm 1.0$ | $40.6 \pm 0.3$ | $48.0 \pm 0.2$ |
| **NADPH** | $53.6 \pm 0.4$ | $39.5 \pm 0.2$ | $47.0 \pm 0.5$ |
| **dUMP** | $54.7 \pm 0.2$ | $45.2 \pm 0.2$ | $48.9 \pm 0.3$ |

## DISCUSSION

Despite its functional importance, nucleotide metabolism has received much less attention than genomics and protein expression studies, although multiple metabolic enzymes involved in these pathways are currently targets of chemotherapy in humans [38]. While the synthesis of most dNTPs occurs in the cytoplasm using the enzyme ribonucleotide reductase, the formation of the DNA-unique base dTMP relies on the concerted action of the TYMS, DHFR and SHMT enzymes forming the dTMP synthesis complex (dTMP-SC). The dTMP-SC was detected in the nucleus as well as in the mitochondria. Defects in dTMP synthesis increase genomic uracil because of replicative incorporation of dUMP instead of dTMP. In addition, the dTMP synthesis pathway is essential for the replication of viral DNA, as shown for human herpesviruses, which may encode for their own TYMS counterpart or upregulate expression of cellular TYMS and DHFR [39,40].

Here we provide evidence that the three proteins involved in *de novo* dTMP synthesis assemble to form the dTMP-SC in the cytoplasm and not only after nuclear translocation, as it was assumed. The choice to use is-PLA, with respect to co-IP, allowed us to obtain novel information on space and time localization of the complexes. The signal of the dTMP-SC complex was expected mainly in the nucleus, but only a slight increase of the nuclear complex was observed when S-phase synchronised A549 cells were assayed. So far, we did not explore other conditions which were reported to increment nuclear complex formation, such as DNA damage, because this was out of the scope of the present work.

The finding that dTMP-SC can assemble in the cytoplasm has many implications, summarised in scheme 2, that will require further investigation. First, in the cytoplasm SHMT1 and DHFR also participate to the folate cycle, whereas TYMS was shown to take part to the mitochondrial *de novo* dTMP synthesis [6], therefore it is likely that formation of the dTMP-SC complex, may affect not only the dTMP pool but the whole one-carbon metabolism. In addition, given that the complex certainly has a very different surface accessibility with respect to the single enzymes, it is probable that its assembly directly affects/controls other post transcriptional modifications (PTM) such as SUMOylation or the ability of these enzymes to bind RNA [41–43].

The dTMP-SC formation may indeed represent a strategy to compartmentalize the enzymes in the cytosol and the effects of stabilization/destabilization of the complex should be investigated as this may represent an innovative and powerful tool to undermine the rewiring of one carbon metabolism that tumour cells use to sustain proliferation. The evidence that the dTMP-SC is abundant in the cytoplasm also suggests that the presence of DNA is not necessary to trigger complex assembly, allowing us to start an *in vitro* characterization of the complex, which is a prerequisite for future structural characterization and rational drug design.

Complex formation was indeed observed by FWB and IP analysis (Fig. 7). Interestingly, since the Chimera-short construct failed to form the complex, it may be speculated that a longer flexible linker is necessary for TYMS and DHFR to reorient with respect to SHMT1. The dissociation constant (Kd) of the Chimera and SHMT1 complex was estimated to be in the low micromolar range (Fig. 8). We also observed a very clear avidity effect, which likely depends on the ability of SHMT1 to act as a bidentate binder. This is not surprising given that the tetrameric assembly of SHMT1 results in a higher symmetry with respect to Chimera that is dimeric. It is therefore possible that one SHMT1 tetramer binds to two Chimera dimers. The tetrameric assembly appears to be crucial in the binding process, since a dimeric variant of SHMT1 did not form the complex. Interestingly, while the minimal catalytic unit of SHMT1 is the dimer [36,44] only in higher organisms SHMT1 is tetrameric [45]. Together with the increased affinity displayed for polyglutamylated-THF [46,47], the present results suggest that the higher oligomeric state of SHMT1 may have evolved to favour novel protein-protein interactions, thus increasing the complexity of the cellular regulation network. The importance of the aggregation state in the control of the interaction between the SHMT isoforms and other binding partners was previously highlighted by us and other groups [36,48–50]. The unique propensity of the SHMT2 isoform to form protein-protein complexes in the dimeric state, together with the evidence that SHMT1 forms the dTMP-SC only when in the tetrameric form, provided in the present work, further underlines that the two isoforms are indeed evolutionary distinct and fine-tuned to achieve different goals in the cell, strikingly optimized to minimize superposition of tasks and interference between their specific cellular roles.

A small amount of the SHMT1-Chimera complex could be identified by western blot after SEC analysis, providing the proof-of-concept necessary for further studies (Fig. 9). Large scale purification of the dTMP-SC complex proved difficult to obtain by gel filtration, mainly due to the presence of aggregates or higher oligomeric state species of Chimera, possibly favoured by the low ionic strength. This result was not unexpected, given that only protein-protein complexes with K_d_ in the low nano molar range can be separated by chromatographic methods. Interestingly, the DSF analysis showed that dUMP had a clear stabilizing effect on Chimera and possibly also on the complex, a finding that might be exploited to further stabilize it and improve the complex purification.

In the present work, the dTMP-SC could be successfully assembled *in vitro* thanks to the Chimeric construct that we designed. Chimera was shown to be fully competent to complete the thymidylate synthesis cycle when mixed with SHMT1 in the presence of NADPH, L-serine and dUMP, after addition of a catalytic amount of THF, with the folate species cycling until total consumption of reducing equivalents (Fig. 6). This result provides a good starting point to further investigate the kinetic properties of the dTMP-SC and to eventually search for possible inhibitors of the entire complex, which may significantly differ from those found for the individual components.

In conclusion, the characterization of the cellular dynamics and structural/functional properties of the dTMP-SC presented here provide a significant advance to understand the role of this complex in normal and cancer cells. At this stage it cannot be excluded that other proteins or binding partners may interact with the dTMP-SC and modulate its assembly, including components of the replicase machinery in the nucleus [10] or other nucleic acids such as RNA in the cytosol. Intriguingly, both TYMS and DHFR bind to their own mRNA in the absence of substrates [43] and we recently demonstrated that also SHMT1 binds RNA, which inhibits the enzymatic activity in a selective way [12,42]. Regulation of the metabolic activity by RNA molecules, was recently found also for enolase 1 [51], strongly suggesting that riboregulation of cellular metabolism may take unexpected routes. Undoubtedly, we are witnessing an era in which an increasing number of entangled regulatory mechanism are emerging as central for the cellular control system: PPI, riboregulation, miRNA, microproteins, are just some examples of interactions that shape a regulatory network that is far more sophisticated than could be predicted. We think that the dTMP-SC assembly is certainly playing a crucial role in this network, especially in cancer cells. Finding the missing component(s) will allow to further characterize the complex and advance in the structural analysis, which is certainly the most challenging goal.

## MATERIAL AND METHODS

### Chemicals and reagents

thymidine (T1895), paraformaldehyde (PFA), sucrose, phosphate buffered saline (PBS), Tween 20, Bovine serum albumin (BSA), 4,6-diamidino-2-phenylindole (DAPI), mounting media, isopropil-β-D-1-tiogalattopiranoside (IPTG), potassium dihydrogen phosphate, di-Potassium hydrogen phosphate, Phenylmethanesulfonyl fluoride (PMSF), Deoxyribonuclease I from bovine pancreas (Dnase), 4-(2-Hydroxyethyl)piperazine-1-ethanesulfonic acid (HEPES), magnesium chloride (MgCl2), sodium chloride (NaCl), Glycerol, Triton X-100, imidazole, 2-mercaptoethanol, 2’-Deoxyuridine 5’-monophosphate disodium salt (dUMP), β-Nicotinamide adenine dinucleotide phosphate (NADPH), L-serine and Duolink PLA kit (DUO92007) were purchased from Sigma-Aldrich. Tetrahydro-folate (THF) was provided from Merck & Cie (Schaffhausen, Switzerland). Protein G-Plus agarose beads (Sc-2002) were purchased from Santa Cruz Biotechnology, INC. Dulbecco’s phosphate buffered saline (DPBS, 20-031-CV), fetal bovine serum (35-015-CV) and 0,25% trypsin (25-053-CI) were purchased from Corning; BCA kit (quantumMicro Protein, EMP015480, EuroClone).

### Antibodies

the following antibodies were used: rabbit anti-SHMT1 (HPA0233314, Atlas antibodies); mouse anti-DHFR (WH0001719M1, Sigma-Aldrich); rabbit anti-TYMS (MBS126074, Mybiosource)indicated as TYMS_R_; mouse anti-TYMS (sc-33679, Santa Cruz Biotechnology, INC) indicated as TYMS_M_; mouse anti-PCNAAlexa Fluor^®^ 488-conjugated antibody (ab201672, Abcam, PC10 clone), mouse anti-α-Tubulin (T5168, Sigma Aldrich, Clone B512); mouse anti-tubulin-FITC (f2168, Sigma Aldrich); rabbit anti-Histone-H3 (H0164, Sigma Aldrich); mouse anti-SHMT1 (sc-365203, Santa Cruz Biotechnology, INC); rabbit anti-SHMT1 (D3B3J, cell signaling technology); anti-DHFR (sc-377091; Santa Cruz Biotechnology, INC); anti-chicken tubulin (ab89984, Abcam); anti-mouse FITC (AB_2338589 Jackson ImmunoResearch); anti-rabbit FITC (AB_2337977, Jackson ImmunoResearch); anti-mouse Rhodamine (AB_2338766 Jackson ImmunoResearch); anti-rabbit CY3 (AB_2338000, Jackson ImmunoResearch); mouse anti-rabbit IgG-HRP (sc-2357, Santa Cruz Biotechnology, INC); m-IgGκ BP-HRP (sc-516102, Santa Cruz Biotechnology, INC); anti-chicken alexa fluor 647 (703605155, Jackson ImmunoResearch); Duolink *is*-pla probe anti-mouse minus (duo92004-100rxn, Sigma Aldrich); Duolink *is*-pla probe anti-rabbit plus (duo92002-100rxn, Sigma Aldrich).

### Cell lines

A549 lung cancer cell lines were purchased from ATCC (Manassas, VA, USA). The cells were grown in RPMI-1640 medium (Corning), supplemented with 100 IU/ml penicillin/streptomycin (P 4458, Sigma Aldrich) and 10% foetal bovine serum (FBS, Corning). HeLa cells purchased from ATCC (CCL-2), cultured in DMEM and supplemented with 2% penicillin/streptomycin, 2% L-glutamine, 2,5% Hepes and 10% foetal bovine serum.

All experiments were run in triplicate and in separate biological sets.

### Cells synchronization

cells were synchronised in S-phase by a single thymidine block. Cells were treated as follows: a) 2mM thymidine (T1895, Sigma-Aldrich) for 24 hours at 37°C, to block DNA synthesis; b) release in a thymidine-free medium; c) after 4 hours cells were fixed in 3.7% paraformaldehyde/30 mM sucrose for 10 min and processed either for the is-PLA experiment by using the Duolink PLA kit (DUO92007, Sigma-Aldrich) according to manufacturer’s instructions, or for the immunofluorescence (IF).

### Immunofluorescence (IF)

IF staining was performed to check whether the cells were synchronised in S-phase (Fig. S1), and to assess the differential cellular localization of the single proteins (SHMT1, TYMS and DHFR) according to the cellular phase, and to co-localize the proteins.

Asynchronous and S-phase synchronised A549 (lung cancer cell lines) were grown on coverslips and fixed in 3.7% paraformaldehyde/30 mM sucrose for 10 min. For the co-localization samples, asynchronous A549 were fixed by using ice cold methanol and incubating the coverslips at −20°C for 6 minutes. Afterwards, cells were permeabilized in 0.1% Triton X-100 and blocked in 3% bovine serum albumin in PBS/0.05% Tween-20 for 1h at room temperature. The incubation with primary antibodies was performed at 4°C o.n. in a dark humidity chamber. Subsequently, the fluorescently labelled secondary antibodies were added in a PBS containing 0.05% Tween 20 and 3% BSA solution and incubated at RT for 30 min. Cells were counterstained with 4,6-diamidino-2-phenylindole (DAPI, 0.1 μg/mL) and mounted by using the mounting media.

Primary antibodies: i) rabbit anti-SHMT1 (1:50), ii) mouse anti-DHFR (1:50), iii) rabbit anti-TYMS (1:20), iv) anti-PCNAAlexa Fluor^®^ 488-conjugated antibody (1:1000), v) anti-tubulin-FITC (1:300), vi) anti-chicken tubulin (1:50), vii) mouse anti-TYMS (1:50).

Secondary antibodies: i) anti-mouse FITC (1:100), ii) anti-rabbit FITC (1:50), iii) anti-mouse Rhodamine (1:50) and iv) anti-rabbit CY3 (1:500) v) anti-chicken alexa fluor 647(1:100).

### *In site*-Proximity ligation assay (is-PLA)

*is*-PLA was performed in both synchronised and asynchronous A549 and HeLa cell lines, by using the Duolink PLA kit and according to manufacturer’s instruction. The primary antibodies pair to detect the interaction among the three proteins were:

i. mouse anti-DHFR (1:50)/ rabbit anti-SHMT1 (1:50)
ii. rabbit anti-TYMS (1:20)/ mouse anti-DHFR (1:50)
iii. rabbit anti-SHMT1 (1:50)/ mouse anti-TYMS (1:50)

The above-mentioned antibodies were incubated o.n. in a dark humidity chamber at 4°C. Subsequently, the incubation with the PLA probes was performed in a pre-heated humidity chamber for 1 hour at 37°C followed by the ligation addition. If the target proteins (SHMT1-DHFR-TYMS) are interacting among each other or are very close, this step will produce a DNA circle. The amplification time was 100 min and was performed at 37°C in a dark humidity chamber. Negative controls: PLA experiment was performed without one of the primary antibodies (Fig. S2). DNA was stained with DAPI as described above.

### Cells fractionation

Together with IF, cell fractionation was performed to analyse the cellular localization of the single proteins (SHMT1, TYMS and DHFR) according to the cellular phase.

Nuclear and cytosolic fractionation of A549 cell lines (both asynchronous and S-phase enriched) was performed by treating with trypsin and washing the cells twice in ice-cold PBS. Cells were lysed in 200 μl of cytoplasmic lysis buffer (Tris Ph 7,8 10mM, MgCl_2_ 1,5 mM, KCl 10mM, 1mM PMSF) and kept on ice for 10 min. Nuclei were then pelleted at 1000xg for 7 minutes at 4°C, and the supernatant containing the cytoplasmic fraction was removed. Nuclei were resuspended in 60 μl of SDS 10% incubated for 30 min on ice, boiled for 10 min and the DNA fraction was sedimented by centrifugation at 1000xg for 5 minutes at 4°C. Protein concentrations were determined by the BCA assay. Cytosolic and nuclear fractions (20 μg) were separated by SDS-PAGE and transferred into a nitrocellulose membrane. Western blot analysis was performed as usual and the primary antibodies diluted 1:1000. WB was performed by using mouse anti-SHMT1 (1:1000, Cell signalling), mouse anti-DHFR (1:1000), rabbit anti-TYMS (1:1000). Anti-tubulin (cytosolic marker, 1:1000) and anti-Histone H3(nuclear marker, 1:5000)were used as controls

### Microscopy analysis on fixed samples

Samples were acquired using a Nikon Eclipse 90i microscope (Nikon Instruments S.p.A., Campi Bisenzio Firenze, IT, USA) equipped with 40× (N.A. 0.75) and and 100x (oil immersion, N.A. 1.3) objectives and a Qicam Fast 1394 CCD camera (QImaging) or with an inverted microscope (Eclipse Ti, Nikon) using a 60× (oil immersion, N.A. 1.4) objective and the Clara camera (ANDOR technology). Images were acquired using NIS-Elements AR 3.2 (Nikon) or Nis-Elements H.C. 5.11 using the JOBS module for automated acquisitions; elaboration and processing was performed using NIS-Elements HC 5.02 (Nikon) and Adobe Photoshop CS 8.0.

### Quantitative analyses of fluorescent signals

For immunofluorescence quantification of fluorescence intensity signals, images of interphase cells of asynchronous and S-enriched populations were acquired using 60x or 40x objectives, along the z axis every 0.4 *μ*m for 5 *μ*m. Signals were measured using Nis Elements H.C. 5.02 (nd2 file format) as follows: sum intensity values of the nuclear fluorescence respect to sum intensity values of the cytoplasmic fluorescence. To avoid differences due to different cell sizes, for each cell the same square mask was used to calculate the nuclear and the cytoplasmic fluorescence. Images were corrected for external background. For is-PLA spots of interaction counts images were acquired with 60x objective, along the z axis every 0.4 *μ*m for 8 *μ*m. The “general analysis” module of Nis Elements H.C. 5.11 was then used for automatic spot count; is-PLA signals were identified based on fixed parameters in all images and those inside the nuclei (defined by DAPI signal) were counted.

Images for quantification fluorescence signals were Maximum Intensity Projections from z-stacks (0.4 *μ*m for a range of 5-8 *μ*m). Statistical analyses were performed with GraphPad Prism 8.

### Chimera rational design

Structural analysis necessary to design and model the two constructs of Chimera was largely conducted within PyMOL [The PyMOL Molecular Graphic System, Version 1.8, Schrödinger, LLC] taking advantage of the PyMod2.0 plugin [52] that allows to perform sequence and structural alignments as well as homology modelling by integrating BLAST, Clustal Omega, MUSCLE, CEalign and MODELLER [53] into PyMOL. The structure of DHRF-TYMS from *B. bovis* (PDB ID 3I3R) was used as a reference to orient the structures of human DHFR and TYMS (PDB ID 1DRF; 1HZW [54,55]) the linker regions were modelled using the *de novo* loop modelling function in MODELLER and the N-terminal portion of hTYMS was modelled based on the closest homologous structure (TYMS from *M. musculus* - PDB ID: 4EB4 [56]). The final Chimera models were further refined using the sculpting function in PyMOL and MOE [Molecular Operating Environment (MOE), 2019.01; Chemical Computing Group ULC, 1010 Sherbooke St. West, Suite #910, Montreal, QC, Canada, H3A 2R7, 2020] using the conjugate gradients method for energy minimization. Surface electrostatic potentials were calculated using APBS (Adaptive Poisson-Boltzmann Solver) [57], also available as PyMOL plugin. Sequence conservation analysis was performed using CAMPO [58].

### Protein expression and purification

*Chimera*: Long and Short Chimera genes were synthesized and optimized for expression in *Escherichia coli* by GeneArt (Life Technologies), cloned into a pET28a expression vector (Invitrogen), using NdeI and XhoI restriction sites. Small scale expression tests were made on two different *Escherichia coli* expression strains varying: isopropil-β-D-1-tiogalattopiranoside (IPTG) concentration; temperature and expression time; lysis protocol (Table 2). The overexpressed fusion proteins were largely present in the inclusion bodies. However, good results, in terms of solubility and yield of protein, were obtained by basal expression of the protein in BL21(DE3) cells, grown at 25° for 24 h, without IPTG, and cell lysis by sonication. This protocol was therefore applied for larger scale expressions of both Chimera-Long and Short as N-terminal His-tagged fused proteins. Cells were harvested by centrifugation at 4°C, resuspended in the lysis buffer (0,1M K-phosphate pH 7,5; 1mM PMSF; 4 mg/mL Dnase; 5mM MgCl_2_; 5% Glycerol) and lysed by ultrasonic disruption on ice. The supernatant, supplemented with 10 mM imidazole, was purified by IMAC on a Ni^2+^-His-Trap column (GE Healthcare, Chicago, IL, USA) in buffer 0,1M K-phosphate pH 7,5, 10 mM imidazole. The IMAC fractions containing the protein were concentrated by ultrafiltration (Amicon, Millipore) and then injected onto a Superdex 200 10/300 FPLC column (GE Healthcare, Chicago, IL, USA) equilibrated with buffer K-phosphate 0,1M pH 7,5 and eluted with a flow rate of 0,9 mL/min. The column was calibrated using the following protein standards; aprotinin, carbonic anhydrase, ovalbumin, conalbumin, aldolase, ferritin (Sigma): R^2^ was 0.994. The purity of the protein was assessed by SDS PAGE. The molar extinction coefficient at 280 nm was determined experimentally by BCA assay (Sigma) according to the manufacturer’s specifications and was 1,51 (1 mg/ml) or 91355 M^−1^cm^−1^. The purified proteins were aliquoted, frozen in liquid nitrogen and stored at −20° C until use. *SHMT1*: was expressed and purified as described previously [50].

**Table 2.**
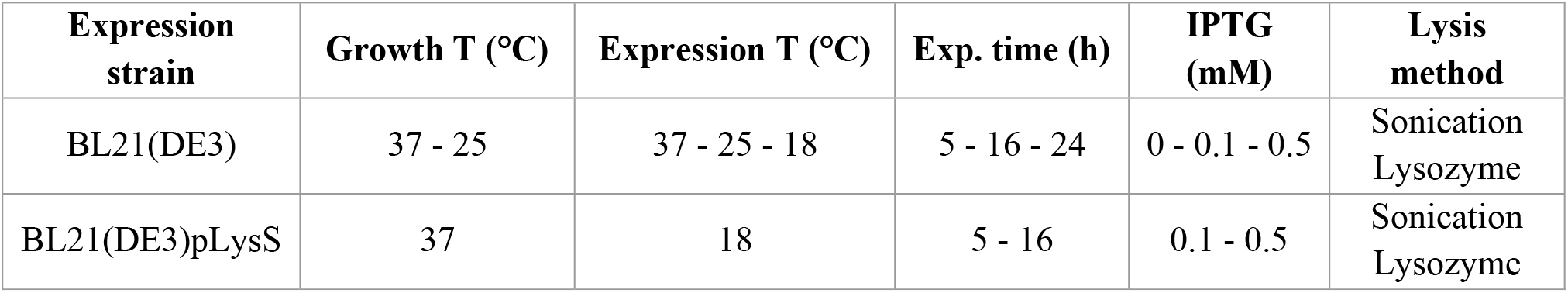
Heterologous expression conditions assayed for Chimera-Short and Long

| **Expression strain** | **Growth T (°C)** | **Expression T (°C)** | **Exp. time (h)** | **IPTG (mM)** | **Lysis method** |
| --- | --- | --- | --- | --- | --- |
| BL21(DE3) | 37 - 25 | 37 - 25 - 18 | 5 - 16 - 24 | 0 - 0.1 - 0.5 | Sonication<br>Lysozyme |
| BL21(DE3)pLysS | 37 | 18 | 5 - 16 | 0.1 - 0.5 | Sonication<br>Lysozyme |

### Circular Dichroism CD and Thermal Melting (TM) spectroscopic analysis

the CD spectra were collected, for both Short and Long Chimera, at a 10 μM protein concentration in a 0.1 cm quartz cuvette (Hellma) using a JASCO J-710 spectropolarimeter equipped with a Peltier temperature control unit. The thermal stability of the two proteins was measured through the TM assay method of the instrument, by increasing the temperature from 30°C to 90 °C with a 1 °C/min rise in T, monitoring the dichroic signal at 220 nm every 0.5 °C.

### Far-Western blot

The prey (Chimera (4 μg), SHMT1 (4 μg)) and the control protein - a truncated construct of RmcA form *P. aeruginosa* (4 μg) [59], SHMT1 or SHMT1-dimeric mutant (H135N-R137A) (0,5 μg), Chimera (0,5 μg) - were resolved by SDS-PAGE electrophoresis, and electro-blotted onto PVDF membranes (Biorad: 100 V, 2h, 4°C). The transferred proteins were then renatured by progressively reducing the guanidine-HCl concentration, starting from the following stock solution: AC buffer (10% glycerol; 5M NaCl; 1M Tris pH 7,5; 0,5 M EDTA; 10% Tween-20; 8M Guanidine-HCl; 2% Milk; 1mM DTT) as described by Wu et al. [60]. Proteins were renaturated by incubating the membranes for 30 min at R.T. in AC buffer containing decreasing concentrations of guanidine-HCl (6M and 3M), then for 30 min at 4°C in 0.1M guanidine-HCl and finally o.n. at 4°C in AC buffer containing no guanidine-HCl. The following day the membranes were blocked with 5% milk in PBS-Tween for 1h at RT and incubated with purified bait proteins (SHMT1 or SHMT1-dimeric mutant and CHIMERA 50 ng/ml) at 4°C overnight. Membranes were then washed and incubated with primary antibody against the protein used for the incubation step – mouse anti-DHFR (1:1000), mouse anti-TYMS (1:1000) and rabbit anti-SHMT1 (1:1000; Cell Signalling) - o.n. at 4°C. Subsequently, membranes were incubated with the appropriate secondary antibody and detected with Luminata™ Crescendo Western HRP Substrate (Millipore) according to manufacturer’s instructions.

### Immunopurification with Protein G-plus agarose beads

SHMT1 and Chimera Long were mixed in a 1:2 ratio in 50 mM NaCl, 20 mM Hepes pH 7.5. Each sample was 1 ml in volume and contained SHMT1 (40 μg); Chimera (92 μg) or the mixed proteins. A precleaning step of the samples was performed by adding 50 μl of Protein G-Plus agarose beads suspension (Santa Cruz Biotechnology, INC) to 1 ml of each sample (Chimera, SHMT1 and Mix) and incubating for 2 h at 8°C on a rocking platform. The beads were then centrifuged at 12.000 xg for 20 s and the supernatant was transferred to fresh tubes. Afterwards, 1 μg of anti-SHMT1 (Santa Cruz Biotechnology, INC) or anti-TYMS (Santa Cruz Biotechnology, INC) was added to each sample, incubating for 1 h at 8°C rocking. 50 μl of beads were then added to the mixture and the final solution was left rocking at 8°C o.n.. Samples were centrifuged at 12.000g for 20 s and the supernatant was discarded. Two washes were performed by resuspending the beads in 500 ul 50 mM NaCl, 20 mM Hepes pH 7,5 and rocking for 30 min at 8°C. The supernatant was discarded. 50 μl of SDS-page loading buffer 2x were added to the beads-protein mixture. Samples were then heated at 95°C for 1h, resolved by SDS-PAGE electrophoresis and subsequently electro-blotted onto a nitrocellulose membrane following a standard western blot protocol. For the detection of the prey proteins mouse anti-DHFR (1:1000), mouse anti-TYMS (1:1000) and rabbit anti-SHMT1 (1:1000; Cell Signalling) antibodies were used.

### Surface Plasmon Resonance (SPR)

SPR experiments were performed at 25 °C using a Pioneer AE optical biosensor (Molecular Devices-ForteBio) equipped with a COOH2 sensor chip (short carboxylated polysaccharide coating), and equilibrated with running buffer 10mM Hepes pH7.4, 150mM NaCl, 0.005% Tween-20. The COOH2 sensor chip was installed and conditioned according to manufacturer’s protocol. Then, the chip was chemically activated for 6 min by injecting 150 μL of a 1:1 mixture of 100 mM N-hydroxysuccinimide (NHS) and 400 mM ethyl-3(3-dimethylamino) propyl carbodiimide (EDC), diluted 1:10 with water, at a flow rate of 25 μl/min. Chimera Long was immobilized onto the activated sensor chip by using standard amine-coupling procedures [61]. Briefly, a 0.1 mg/mL Chimera solution (in 10 mM sodium acetate, pH 4.5) was injected at a flow rate of 10 μL/min on channels 1 and 3 (channel 2 was used as reference for nonspecific binding), followed by a 70 μL injection of 1 M ethanolamine pH 8.0 to block any remaining activated groups on the surface. Chimera was captured to a density of 735 and 175 resonance units (RUs) on Ch1 and Ch3 channels, respectively. The stability of the Chimera surface was demonstrated by the flat baseline achieved at the beginning (0–60 s) of each sensorgram. The analyte SHMT1 was dialyzed in the running buffer and injected at different concentrations (85, 42.5, 21.2,10.6, 5.3, 2.66 μM) onto the sensor chip at a constant flow rate of 75 μl/min. A dissociation of 180 s was allowed. At least two experiments were performed. The interaction of the analyte with immobilized Chimera was detected as a measure of the change in mass concentration, expressed in RUs. All sensorgrams were processed by using double referencing [62]. First, responses from the reference surface (Ch2) were subtracted from the binding responses collected over the reaction surfaces to correct for bulk refractive index changes between flow buffer and analyte sample. Second, the response from an average of the blank injections (zero analyte concentrations) was subtracted to compensate for drift and small differences between the active channel and the reference flow cell Ch2 [63]. To obtain kinetic rate constants and affinity constants the corrected response data were fitted in the QDAT software. A kinetic analysis of the ligand/analyte interaction was obtained by fitting the response data to a reversible 1:1 bimolecular interaction model. The equilibrium dissociation constant (k_d_) was determined by the ratio k_off_/k_on_. This experiment was performed in duplicate.

### Chromatography

the formation of dTMP-SC complex between SHMT1 and Chimera Long was analysed by HPLC-size exclusion chromatography and compared to the individual proteins. SHMT1 and Chimera were mixed in a 1:2, 1:1 or 2:1 molar ratio, with 1 corresponding to a final concentration of 35 μM (monomer) for both proteins in the 1:1 sample and to 17.5 μM in the 1:2 and 2:1. All samples were incubated at 4°C overnight prior to the analysis. 100 μl of each sample was injected onto an HPLC column (TSK G3000PWXL 7,8 mm x 30 cm with a 4 cm precolumn; TOSOH bioscience) equilibrated with 20 mM potassium phosphate pH 7.5, at a flow rate of 0.7 ml/min, and detected at 280 nm. Fractions corresponding to the putative complex were collected, concentrated and analysed by SDS-PAGE and WB (rabbit anti-SHMT1, 1:1000; Cell signalling). The column was calibrated using the following protein standards (Sigma); ferritin (12 nm – 440 KDa); aldolase (11 nm – 158 kDa); conalbumin (8.8 nm – 75 KDa); ribonuclease A (6.7 nm 13.7 KDa) - R2 of the linear fit was 0.995. The column was connected to an HPLC pump (Labflow4000 - Labservice analytica, Italy) equipped with an UV detector (Smartline 2520 - Knauer, Germany).

### Differential Scanning Fluorimetry (DSF)

DSF assays were performed using a Real Time PCR Instrument (CFX Connect Real Time PCR system, Bio-Rad). In a typical experiment, 0.5 μM SHMT1 and 0.5 μM Chimera (Long or Short) in 20 mM Na HEPES pH 7.2, containing 50 mM NaCl, were incubated o.n. at 4°C (in a total volume of 25 μl) in a 96-well PCR plate. When substrates were added, this solution also contained 0.1 mM CHO-THF, 1 mM dUMP, 0.1 mM NADPH. After the incubation, Sypro Orange (4x, Thermo Scientific) was added and fluorescence was measured from 25°C to 95°C in 0.4°C/30 sec steps (excitation 450-490 nm; detection 560-580 nm). All samples were run in triplicate. Denaturation profiles were analysed using the PRISM Graph Pad software, after removal of points representing quenching of the fluorescent signal due to post-peak aggregation of protein:dye complexes. All curves were normalised and fitted to the following sigmoidal equation to obtain the melting temperatures (T_m_).

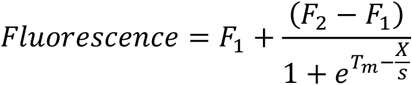

where X is the temperature (°C), F1 is the fluorescence at low temperature and F2 the maximal fluorescence at the top of the truncated dataset, while s describes the steepness of the curves. Alternatively, T_m_ values were obtained by plotting the first derivative of the fluorescence emission as a function of temperature (-dF/dT) by using the Bio Rad CFX manager software.

### Activity assays

activity assays were carried out at 20°C in 20 mM K-phosphate pH 7.2, containing 75 mM 2-mercaptoethanol, using a Hewlett-Packard 8453 diode-array spectrophotometer (Agilent Technologies, Santa Clara, CA) and a 1-cm pathlength cuvette. Concerning TYMS activity assay, substrate concentration was 0.2 mM CH_2_-THF, 0.1 mM dUMP and varying concentration of Chimera (0.3, 0.6 and 1.1 μM). Saturation curves, from which TYMS kinetic parameters were calculated, were obtained using 1 μM Chimera keeping one substrate at fixed concentration (0.1 mM dUMP or 0.05 mM CH_2_-THF) and varying CH_2_-THF from 0 to 0.2 mM or dUMP from 0 to 0.1 mM. In order to calculate kinetic parameters, the saturation curve shown in Fig. 9C left panel was fitted to an equation describing the substrate inhibition [24], whereas the saturation curves in Fig. 9C right panel was fitted to the Michaelis-Menten equation. Concerning TYMS-DHFR activity, substrates concentration was 0.1 mM CH_2_-THF, 0.1 mM dUMP, 0.1 mM NADPH, whereas Chimera was 0.5 μM. In the activity assay of dTMP-SC substrates concentration was 10 mM L-serine and 16 μM THF, 0.1 mM dUMP and 0.1 mM NADPH. Chimera and SHMT1 concentration was 0.5 μM.

## AUTHOR CONTRIBUTIONS

SS, APai, DB, GB, GP, Apao, SR, AT, GG, FC: planned experiments; SS, DB, GB, RL, FRL, DC, RM, GP, AT, GG: performed experiments. SS, DB, RL, GP, APao, SR, RC, APai, AT, GG, FC: analysed data. SS, DB, RL, GP, AT, GG, FC wrote the paper. All the authors critically revised and approved the final version of the manuscript.

## ACKNOWLEDGMENTS

Imaging experiments were performed at the Nikon Reference Centre, CNR Institute of Molecular Biology and Pathology.

## FUNDING

This work was supported by Associazione Italiana Ricerca sul Cancro (AIRC) [IG 2019 - ID. 23125] P.I. F.C.; and Sapienza University of Rome [RG11816430AF48E1, RM11916B46D48441, RP11715C644A5CCE, GA116154C8A94E3D-HypACB platform] to F.C.

## ABBREVIATIONS

dTMP: Deoxythymidine monophosphate
dUMP: deoxyuridine monophosphate
TYMS: thymidylate synthase
SHMT: serine hydroxymethyltransferase
DHFR: dihydrofolate reductase
CH_2_-THF: 5,10-methylene-tetrahydrofolate
THF: tetrahydrofolate
DHF: dihydrofolate
dTMP-SC: thymidylate synthesis complex
IF: Immunofluorescence
is-PLA: *in situ* proximity ligation assay
IPTG: isopropil-β-D-1-tiogalattopiranoside
5-FU: 5-fluorouracil
MTX: methotrexate
PTX: pemetrexed
FdUMP: fluorodeoxyuridine-monophosphate
cryo-EM: cryogenic electron microscopy
IMAC: immobilized metal affinity chromatography
DFS: Differential scanning fluorimetry
SPR: surface plasmon resonance
o.n.: overnight
R.T.: room temperature.

## SCHEMAS

**Scheme 1.**
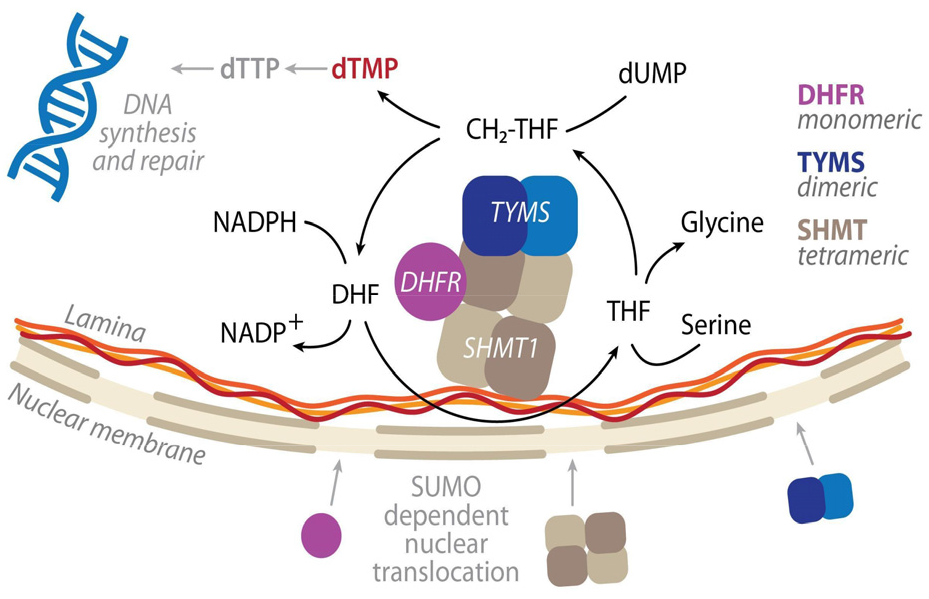
Scheme of the nuclear thymidylate synthesis complex (dTMP-SC) catalytic cycle. SHMT, DHFR and TYMS are SUMOylated and translocate to the nucleus during G1/S-phase, where they are proposed to assemble to form the dTMP synthesis complex (dTMP-SC), anchored to the nuclear lamina [7]. The oligomeric state of the three enzymes is also reported.

**Scheme 2:**
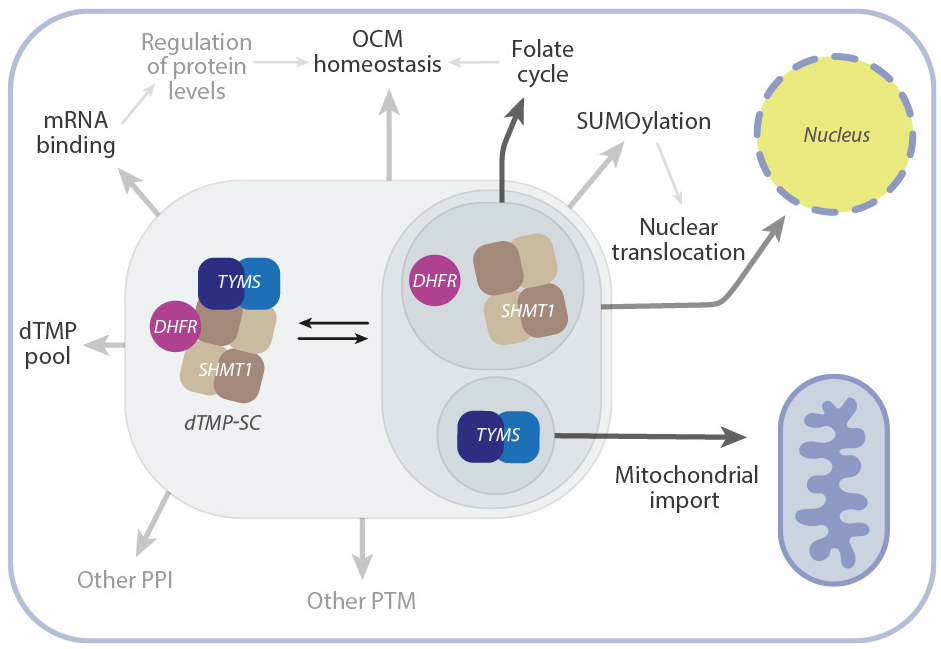
dTMP-SC formation equilibrium in the cytosol. Scheme of the processes that may be affected by the assembly of the dTMP-SC in the cytosol. After complex formation, some regions of the three enzymes may become less accessible. This may in turn affect: i) the SUMOylation/deSUMOylation equilibrium which controls nuclear translocation and possibly other functions; ii) the ability of the proteins to bind mRNA and cross regulate protein homeostasis and, in the case of SHMT1, also the catalytic activity (riboregulation) [42] with a direct effect on the folate cycle and OCM; iii) other PPI or PTM as well as the mitochondrial import of TYMS

## Notes

### Competing Interest Statement

The authors have declared no competing interest.

### Summary of Updates

Addition of Co-localization experiments. Figures update to include all supplementary data that are now no longer needed.

